# Primary human colonic mucosal barrier crosstalk with super oxygen-sensitive *Faecalibacterium prausnitzii* in continuous culture

**DOI:** 10.1101/2020.07.02.185082

**Authors:** Jianbo Zhang, Yu-Ja Huang, Jun-Young Yoon, John Kemmitt, Charles Wright, Kirsten Schneider, Pierre Sphabmixay, Victor Hernandez-Gordillo, Steven J. Holcomb, Brij Bhushan, Gar Rohatgi, Kyle Benton, David Carpenter, Jemila C. Kester, George Eng, David T. Breault, Omer Yilmaz, Mao Taketani, Christopher A. Voigt, Rebecca L. Carrier, David L. Trumper, Linda G. Griffith

## Abstract

The gut microbiome plays an important role in human health and disease. Gnotobiotic animal and *in vitro* cell-based models provide some informative insights into mechanistic crosstalk. However, there is no existing system for a chronic co-culture of a human colonic mucosal barrier with super oxygen-sensitive commensal microbes, hindering the study of human-microbe interactions in a controlled manner. Here, we investigated the effects of an abundant super oxygen-sensitive commensal anaerobe, *Faecalibacterium prausnitzii*, on a primary human mucosal barrier using a Gut-MIcrobiome (GuMI) physiome platform that we designed and fabricated. Chronic continuous co-culture of *F. prausnitzii* for two days with colon epithelia, enabled by continuous flow of completely anoxic apical media and aerobic basal media, resulted in a strictly anaerobic apical environment fostering growth of and butyrate production by *F. prausnitzii*, while maintaining a stable colon epithelial barrier. We identified elevated differentiation and hypoxia-responsive genes and pathways in the platform compared with conventional aerobic static culture of the colon epithelia, attributable to a combination of anaerobic environment and continuous medium replenishment. Furthermore, we demonstrated anti-inflammatory effects of *F. prausnitzii* through HDAC and the TLR-NFKB axis. Finally, we identified that butyrate largely contributes to the anti-inflammatory effects by downregulating TLR3 and TLR4. Our results are consistent with some clinical observations regarding *F. prausnitzii*, thus motivating further studies employing this platform with more complex engineered colon tissues for understanding the interaction between the human colonic mucosal barrier and microbiota, pathogens, or engineered bacteria.

## Introduction

The gut microbiome has emerged as a key factor regulating and responding to human health and disease. Altered gut microbiome composition has been linked to numerous diseases including autoimmunity, inflammatory bowel diseases (IBD), neurodevelopmental disease, metabolic disorders, cancer, and more recently behavior learning.^1-8^ For example, reduction of a major member of the phylum Firmicutes, *Faecalibacterium prausnitzii*, is strongly associated with a higher risk of ileal Crohn’s disease.^9^ In a chemically-induced colitis mouse model, *F. prausnitzii* and its supernatant markedly alleviated the severity of the colitis.^9^ In a mouse model of autism, neurodevelopmental symptoms have been demonstrated to coincide with microbiota alterations and a “leaky gut”. *Bacteroides fragilis* was demonstrated to ameliorate gut permeability as well as communicative and sensorimotor behavioral defects in this animal model.^10^

While studies in mice provide some informative insights into the mechanisms by which *F. prausnitzii, B. fragilis* and other organisms exert their beneficial effects, they also have limitations: inconsistent translation to human physiology; limited ability to control microbe-gut interactions; and relatively low throughput. Moreover, many bacterial species implicated in human health are highly specific to the human host and do not appear in common mice fecal microbiota; nor do humans host many of the microbes found naturally in mice.^11^ Furthermore, microbes colonize the gut in a microenvironment-specific manner related to nutrient load, mucus properties, and oxygen tension. For example, *F. prausnitzii* is a super oxygen-sensitive anaerobe^12^ (i.e., it is on the extreme end of oxygen intolerance among “obligate anaerobes”) that cannot colonize the microaerobic environment ^13^ of the upper gastrointestinal (GI) tract, but robustly colonizes the anaerobic environment of the human large intestine.^14^ However, these microenvironments are difficult to control and study in animal models. These concerns, together with an increasing emphasis on replacement, reduction, and refinement of animal use^15^ motivate the development of *in vitro* microphysiological systems (MPSs) representing the unique environment of the human colon to enable studies of human-microbe interactions in a controlled manner.

Among the organisms significantly implicated in human health and disease, *F. prausnitzii* is particularly interesting for *in vitro* mechanistic studies, as it is so fastidious that it fails to colonize gnotobiotic mice unless the mice are first colonized with a commensal such as *B. thetaiotaomicron*, a more oxygen-tolerant organism than *F. prausnitzii*;^12,16^ further, sustained colonization typically requires repeated inoculations.^17^ In humans, the relative deficiency of *F. prausnitzii* in both UC and CD patients concurrent with a robust increase in TLR4 observed in the intestinal epithelial cells of these patient populations^18^ suggests a possible role for *F. prausnitzii* in modulation of TLRs in the human intestine. TLR receptors are critical for intestinal recognition of bacteria, but activation of TLRs is also associated with an increase of NFKB signaling and inflammation. Imbalanced relationships within this triad may promote aberrant TLR signaling, contributing to acute and chronic intestinal inflammatory processes in IBD, colitis and associated cancer.^19,20^ The responsiveness of intestinal cells to LPS is positively correlated with TLR4 expression.^21^ Thus reduction of factors that keep TLR expression in check– potentially, *F. prausnitzii* –may contribute to chronic intestinal inflammatory diseases. *In vitro* studies to test hypotheses about *F. prausnitzii-*mediated regulation of colon epithelial function or immune functions have been limited to <12-hr static culture by technical difficulties in maintaining an anaerobic apical environment supporting viable microbes in the context of an aerobic basal environment to maintain epithelial cell integrity.^22-24^ As appreciated by these previous investigators, the influences of *F. prausnitzii* on cellular phenotype may arise from relatively long (>24 hr) time scale phenomena, such as accumulation of butyrate and other microbial metabolites,^25^ in addition to other transcriptionally-controlled behaviors. Recent technological innovations to culture obligate anaerobes in modified Transwell formats are restricted to static cultures – where microbial nutrition become limiting – and have so far been used to culture anaerobes that are not as strictly oxygen sensitive as *F. prausnitzii*.^26,27^

In order to study the long-term effects of live *F. prausnitzii* on human primary colon epithelial barriers, we engineered a microfluidic platform to allow chronic (days) co-culture of super oxygen-sensitive microbes with a primary human colon epithelial barrier maintained on a standard cell culture membrane insert (Transwell). Recognizing that maintenance of a robust microbial population for long culture periods requires frequent refreshment of the microbial culture medium, a crucial platform feature is programmable apical fluid flow, which can provide a continuous flow of completely anoxic media at a desired rate. Further, to enable quantitative analysis of steroids and lipophilic drugs, the platform is constructed from materials that inhibit adsorption of lipophilic compounds. Several continuous microfluidic devices that meet some of these criteria have been reported in the literature,^28-30^ and though some have supported culture of some species of obligate anaerobes, ^30,31^ none have reported culture of *F. prausnitzii* and it is unclear whether these devices support microbes that fall into the most super oxygen-sensitive range. Further, most microfluidic devices are fabricated from polydimethylsiloxane (PDMS), which is highly adsorptive of lipophilic compounds and oxygen permeable.^29,30,32^ Approaches to mitigate the high oxygen permeability of PDMS include placement of devices in a custom anaerobic chamber ^29^ and increasing the thickness of the PDMS. ^30^

Here, we describe the design and implementation Gut-MIcrobiome (GuMI) mesofluidic culture platform for analysis of interactions between *F. prausnitzii* and a primary human colon mucosal barrier over a 4-day period. We first examined the effects of platform culture on the phenotype of the colon mucosal barrier compared to traditional static culture in the absence of bacteria, assessing morphological criteria, barrier function, and distribution of cell types via immunostaining, along with analysis of apical cytokine and growth factor accumulation and RNA sequencing analysis for transcriptional regulation. We then studied the behavior of *F. prausnitzii* in multi-day co-culture with the human primary colon mucosal barrier, measuring microbial localization, growth kinetics, and short chain fatty acid (SCFA) production rates. In addition, using RNA sequencing, pathway analyses, and quantitative PCR, we revealed specific responses of colon epithelium to luminal hypoxia and the continuous growth of and butyrate production by *F. prausnitzii*. Expression of some critical autocrine growth factor and cytokine genes GRO, FGF2, TNFa, IL-15, IL-18, and IL-33 were altered in colon epithelium in response to *F. prausnitzii*. In addition, extended co-culture with *F. prausnitzii* was associated with downregulation of toll-like receptor 3 (TLR3) and 4 (TLR4) in the epithelia, consistent with expressional changes strongly associated with IBD in humans.^18^ Downstream signaling pathways NFKB1 and NFKB-activating pathway were also downregulated concomitant with upregulation of the NFKB-inhibitory pathway. We further parsed these *F. prausnitzii*-mediated changes to identify those attributable primarily to its production of butyrate.

## Results

### GuMI physiome platform design and fabrication

Transwell® and related commercially-available membrane culture inserts remain widely used for building *in vitro* models to study epithelial barriers, including gut, lung, cervix and endometrium, as they foster robust monolayer formation, are highly reproducible, and easy to use.^33-36^ We therefore designed the GuMI device to modulate the apical and basolateral microenvironments of these standardized membrane culture inserts individually, by controlling the flow rates and oxygen concentrations separately on the apical and basolateral sides to establish a steep oxygen gradient across the epithelial barrier resembling that in the colon mucosa.^37^

The GuMI physiome platform, designed to maintain six cultures with controlled microenvironments, comprises separate apical and basolateral modules (**Figure 1**). Each module is assembled in three layers: a fluidic plate, a flexible membrane, and a pneumatic plate. Since the fluidic plate is machined from a monolithic block of polysulfone, in contrast to oxygen-permeable PDMS commonly used in microfluidic devices, GuMI is designed to operate in a standard cell culture incubator. The polysulfone plate is both oxygen impermeable and sterilizable through autoclaving. Microfluidic channels and pumps are machined at the bottom of the fluidic plate, where each pump comprises three micro-chambers in line with a fluidic channel. Each chamber can be set to open or close by applying either vacuum or pressure to the pneumatic line to flex the membrane located in between the fluidic and pneumatic plates. By operating the chambers in a specific sequence and frequency, the fluidic direction and flow rate can be precisely controlled as described previously.^38,39^ Driving flow with pneumatic control provides a great advantage in parallelism and scalability, and it also allows GuMI to process multiple samples without additional pneumatic lines.

**Figure 1.**
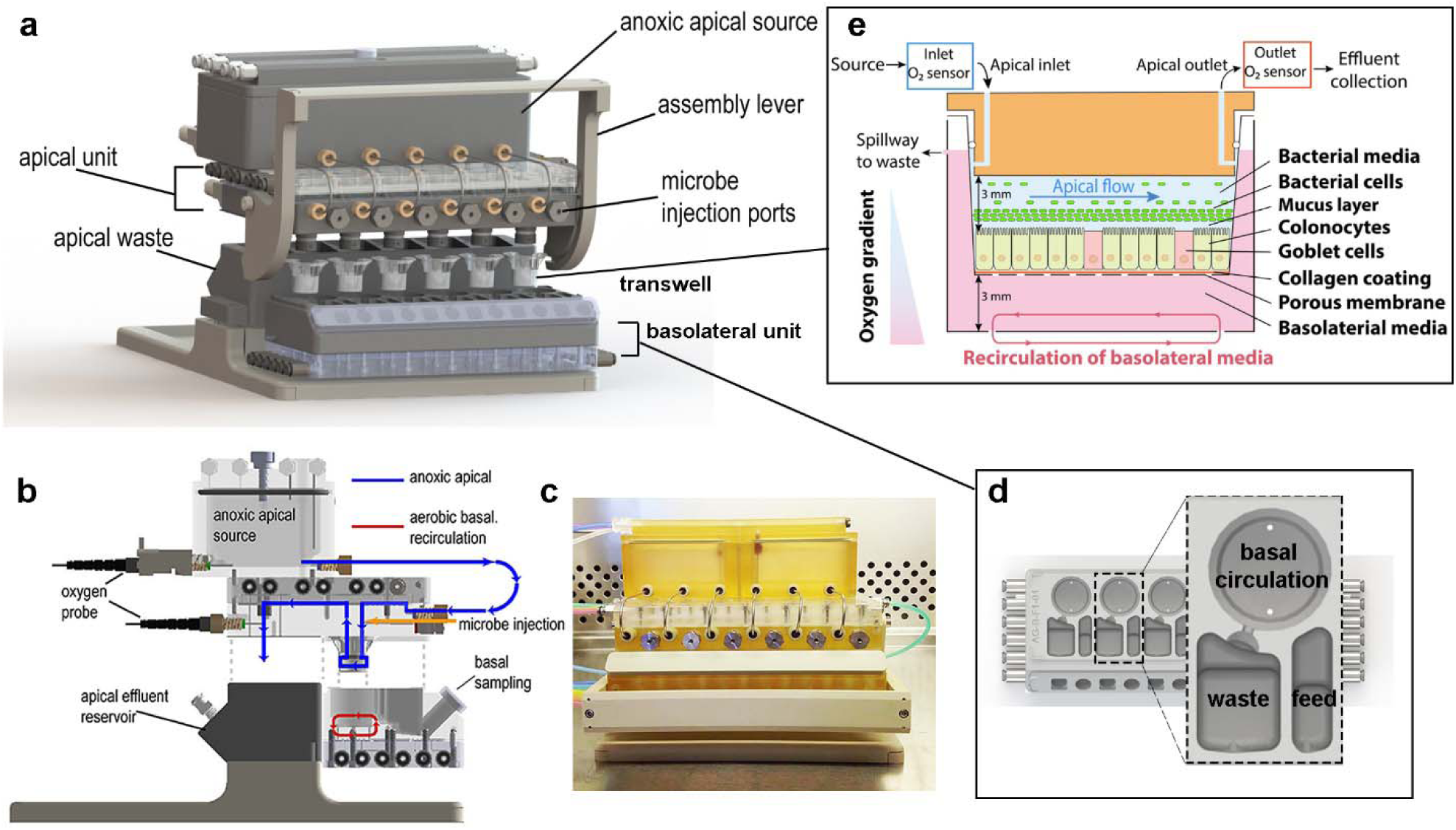
Schematic illustration of GuMI physiome platform. (**a**) The GuMI physiome platform recapitulates a steep oxygen gradient across intestinal epithelium cultured in standardized Transwells to support the co-culture of the intestinal epithelium and obligate anaerobes. The platform is composed of an apical and basolateral unit with individual flow control and is capable of supporting six samples at once. Individual microbe injection ports capped with an oxygen impermeable septum allows obligate anaerobes of interest to be introduced into each sample independently. (**b**) Side illustration of the GuMI physiome platform. Anoxic apical media purged with inert gas is delivered into each Transwell by pneumatic pumps located in the apical units, where the effluent is collected at the apical effluent reservoir. Meanwhile, continuous recirculation of basolateral media by pneumatic pumps inside the basolateral unit promotes oxygenation and supply the oxygen needed for epithelial cells. Oxygen probes at the apical medial reservoir and near the apical effluent exit measure oxygen concentration at apical inlet and outlet respectively. (**c**) Photo of GuMI physiome platform assembled in a sterile cell culture hood. (**d**) Schematic illustration of the basolateral unit of the GuMI platform. Each basolateral unit contains six replicates of a basolateral module. A Transwell is located inside the circular well at the top of the module, with a set of pneumatic pumps underneath providing oxygen through circulation. Another set of pneumatic pumps located underneath the feeding compartment supply fresh media to the basolateral side of the sample, where the spent media gets displaced to the waste compartment via the spillway. (**e**) Schematic illustration of a Transwell located inside the GuMI physiome platform. Differentiated colonic epithelial cells seeded inside a Transwell are cultured inside the GuMI platform to study host-microbe interaction *in vitro*.

In order to keep the apical environment anoxic, the apical medium was first purged with a gas mixture of 5% CO_2_ and 95% N_2_ to remove any dissolved oxygen in the apical medium. Then the medium was drawn into the apical module by the pneumatic pump via stainless steel tubing (**Figure 1b**). To ensure the apical medium stays anoxic in the module, pump chambers were actuated with nitrogen and vacuum, and an additional stream of nitrogen was applied to the space between the membrane and the pneumatic plate to displace any remaining oxygen in the device. An assembly lever located on the apical module is designed to mechanically hold the apical and basolateral modules together (**Figure 1c**). Upon activation of the lever, the compression force helps form a tight seal between the O-ring and the sidewall inside a Transwell, allowing apical effluent to be collected at the apical effluent reservoir (**Figure 1b**).

In vivo, oxygen and nutrients supporting the metabolism of cells are delivered to the colon epithelial cells from the submucosa (basolateral side).^37^ Therefore, a recirculation pump is included in the bottom of the basolateral module to enhance oxygen exchange between the basolateral medium and air inside the incubator. The basolateral module also contains a feed compartment with an additional set of pneumatic pumps to supply fresh medium to the basolateral compartment in a programmable fashion, where the spent medium is spilled and collected at the waste compartment through the spillway design (**Figure 1d**).

The continuous perfusion of anoxic apical medium and recirculation of aerobic basolateral medium maintains a physiologic hypoxia with a steep oxygen gradient across the epithelial layer as shown in **Figure 1e**. Ruthenium-based optical oxygen sensors are incorporated near the apical inlet and outlet for real-time monitoring of oxygen concentration. Once an anoxic condition is reached and maintained, obligate anaerobes grown in the log phase are injected into the lumen via the septum located at the front of GuMI (**Figure 1b** and **1e**).

### Prediction and validation of oxygen tensions

After conducting pilot experiments using Caco-2 monolayers^40^ to establish that the GuMI device operated as designed, i.e., maintained a viable mucosal barrier with a steep oxygen gradient between the aerobic basolateral compartment and the anoxic apical compartment (data not shown), we focused on using primary human colon epithelial cells. Epithelial cells obtained from a healthy region of a colon biopsy were expanded as intestinal organoids and seeded as monolayers in Transwells prior to each experiment following the protocol described earlier.^41^ Briefly, primary epithelial cells seeded in a Transwell at 268,000 cells per cm^2^ (300,000 per well) were allowed to reach confluency in seeding medium for three days, then the cells were switched to differentiation medium for four days before being transferred to GuMI for experiments **(Figure 2a)**. Inside the GuMI platform, the apical compartment was perfused with anoxic apical medium, where the oxygen level was constantly measured at one-minute intervals using probes positioned at the inlet and outlet of the apical compartment during operation with mucosal barrier-microbe cultures. Computational simulation suggested that the apical compartment would reach equilibrium at a completely anoxic state after ∼2.5 h (**Figure S1a** and **S1b**) using parameters in **Table S1**. Experimentally, we observed that for an oxygen level of 0 kPa in the inlet reservoir, the outlet concentration measured at the point of the drip tube – a location that incurs some back-diffusion of oxygen from the outlet hence reflects approach to homeostasis but not exact concentrations in the Transwell region - reached a steady state at around 3 kPa (31 µmol/L, 3% oxygen saturation, 15% air saturation) after 24 hours of perfusion at 10 μl min^-1^ across three different samples as shown by the closely overlapping oxygen measurements (**Figure 2b**). True oxygen concentrations in the Transwell region measured invasively (non-sterile sampling) in pilot experiments were close to zero, and the accuracy of this reading is supported by the microbial culture data described below. Further, the more extended time to approach homeostasis than predicted is likely due to the almost unavoidable appearance of small bubbles in the apical region when placing the Transwell (see the quantitative estimation of bubble purge times in Methods). At this flow rate of 10 μl min^-1^ the shear stress that the monolayer experienced is up to 11 µPa, with the highest shear stress at the inlet and outlet ports (**Figure S1c** and **S1d**). This agrees with the flow distribution pattern across the monolayer (**Figure S1e** and **S1f**). This very low shear stress for a relatively high volumetric flow rate is achieved by a meso-scale height of the upper chamber, 3 mm, compared to the microscale of typical microfluidic culture devices (∼ 0.15 mm high).^32^ The flow rate (0.7-3 ml/min) in the small intestine with a diameter of 2 cm generates average shear stress of ∼20 µPa.^42^ Oxygen concentrations measured in the constantly-mixed basal compartment during microbial-mucosal barrier culture during pilot experiments to establish parameters were typically ∼16 kPa.

**Figure 2.**
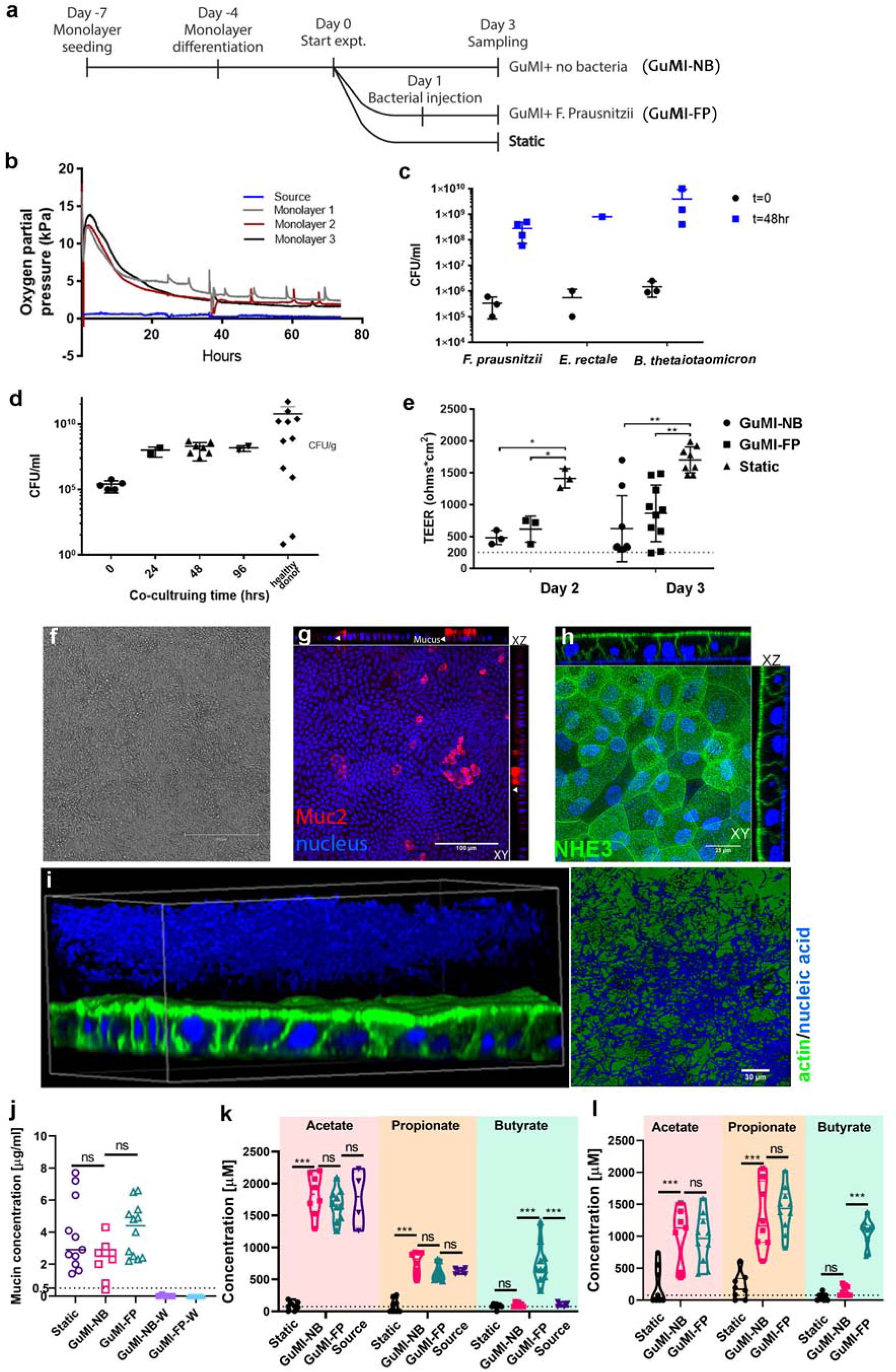
Characterizations of colon epithelial monolayers and their co-culture with obligate anaerobes in GuMI physiome platform. (**a**) Timeline overview of experimental design. For each experiment, monolayers are treated in three conditions: anoxic apical flow (GuMI-NB), anoxic apical flow with *F. prausnitzii* (GuMI-FP) and normoxic static culture (static). (**b**) Representative oxygen measurements of monolayers cultured in GuMI physiome platform. Oxygen concentration is measured at source and near the apical effluent exit for each monolayer to monitor oxygen concentration on the apical side. (**c**) Initial and final concentration (48 hours) of three species of obligate anaerobes (*F. prausnitzii, Eubacterium rectale, B. thetaiotaomicron*) co-cultured with colon epithelium inside GuMI physiome platform. Data are presented as mean ± standard deviation (SD). CFU: colony-forming unit. (**d**) The concentration of *F. prausnitzii* cells co-cultured with colon epithelium inside GuMI at different time points compared to values reported *in vivo*. Data are presented as mean ± standard deviation (SD). CFU: colony-forming unit. (**e**) Transepithelial electrical resistance (TEER) measurements of monolayers on day 2 and 3. GuMI-NB and GuMI-FP have comparable TEER values and are both significantly higher than 300 ohms · cm^2^ (dotted line), suggesting epithelial barriers are intact. Monolayers cultured in GuMI-NB and GuMI-FP have significantly (p<0.05) lower TEER values compared to Static on both day 2 and 3. Data are presented as mean ± SD. (**f-h**) Phase contrast and confocal immunofluorescent staining of monolayers cultured in GuMI-NB for three days. The monolayer is intact and functional and stained positively for different epithelial cell types (**g-h**) with proper apical polarization of NHE3 (**h**). (**i**) Bacterial cells (*F. prausnitzii*, blue) located on top of the colon epithelia (green/blue) from GuMI-FP in 3D reconstruction and 2D top-down view. Green: actin filament staining, blue: 4’,6-diamidino-2-phenylindole (DAPI) nuclei/nucleic acid staining. A ∼15 µm gap in DAPI staining is apparent between the colon epithelia and the zone positive for bacteria. This gap zone likely corresponds to the firm inner mucus layer observed *in vivo* in a mouse distal colon.^48^ (**j**) Concentration of mucin (µg/ml) in the apical media collected directly above the monolayer (GuMI-NB, GuMI FP, and Static) or at the effluent waste (GuMI-NB-W and GuMI-FP-W). Concentration of acetate, propionate, and butyrate in apical side (**k**) and basolateral side (**l**). Data are presented as mean ± SD. Dotted line in (j-k) indicates the limit of quantification. Two-way ANOVA with Dunnett’s multiple comparisons test was used in (k-l). One-way ANOVA with Dunn’s multiple comparisons test was used for Static, GuMI-NB, and GuMI-FP in (e) and (j). ns: not significant; *: p<0.05; ** p <0.01; ***: p<0.001.

### Obligate anaerobe co-culture with colon epithelia in GuMI

The most rigorous test of oxygen levels at the apical surface of the mucosal barrier is maintenance and/or growth of super oxygen-sensitive anaerobes in the apical microenvironment. Therefore, to validate the anoxic apical environment that GuMI is designed to accomplish, three species of obligate anaerobes with varying degrees of oxygen sensitivity (*F. prausnitzii, E. rectale* and *B. thetaiotaomicron*, on the order of most oxygen-sensitive to least oxygen-sensitive), each with significant relevance to human health, were selected to be cultured with colon monolayers independently inside GuMI (**Figure 2c**). The experiments were designed to test the robustness of the platform in supporting oxygen-sensitive anaerobes, thereby establishing the system as an *in vitro* mimic of gnotobiotic conditions useful for mechanistic studies. *F. prausnitzii*, in particular, is characterized as an extremely oxygen-sensitive species, and is strongly implicated in inflammatory bowel diseases;^43-45^ however, mechanistic insights of its effects on human colon remain lacking partly due to its extreme sensitivity to oxygen.^12,46^

*F. prausnitzii, E. rectale* and *B. thetaiotaomicron* at 1-10 × 10^5^ colony forming unit per ml (CFU ml^-1^) were each injected into the apical compartments of separate GuMI Transwell cultures following 24 hours of anoxic perfusion. The microbes were then co-cultured on the apical side of the colon epithelial cells for 2-4 days using standard YCFA bacterial culture medium diluted to 10% (v/v) with phosphate buffered saline flowing continuously at 10 μl/min. The final density of the selected anaerobes measured at the end of the experiment demonstrates that GuMI is capable of supporting the growth of even the most oxygen-sensitive species in isolation, without the support of any facultative anaerobes. Final densities of *F. prausnitzii, E. rectale* and *B. thetaiotaomicron* all reached ∼10^9^ CFU ml^-1^ after 48 hours of co-culturing (**Figure 2c**). This is in stark contrast with control cultures of *F. prausnitzii* maintained in co-culture with a deliberately perforated (leaky to oxygen) epithelial monolayer in GuMI, static Transwell co-culture in the standard incubator environment, or alone under standard static microbial anaerobic conditions. In these cases, the density of microbes was undetectable due to either the presence of oxygen (in co-culture with leaky epithelia or epithelia in static culture in a standard incubator; data not shown) or the lack of nutrients as a result of overgrowth (standard microbial anaerobic culture). A similar trend was observed for *E. rectale*, but *B. thetaiotaomicron* tolerated standard incubator co-culture with colon epithelia, overgrowing and killing the epithelial monolayer within 24 h.

We then focused on the most oxygen-sensitive microbe, *F. prausnitzii*, and determined the growth properties at finer temporal resolution. The bacterial population reached 10^9^ CFU ml^-1^ within 24 hours of inoculation into the apical compartment and remained stable at this concentration thereafter **(Figure 2d)**. This density is consistent with the density observed in healthy human^47^ and mice colon or feces associated with *F. prausnitzii*.^14^ The washout of viable bacterial cells was not monitored as they cannot survive in the effluent reservoir connecting to the incubator atmosphere; however, we presume that the stable concentration reflects a dynamic equilibrium with continued growth accompanied by washout.

### Effects of *F. prausnitzii* on the phenotype of colon epithelial monolayers

After showing the GuMI device supported the growth of *F. prausnitzii* on the apical side of colon epithelial cells cultured in Transwells, we examined the effects of *F. prausnitzii* on the functions of the colon epithelium. One of the most important functions of colon epithelium is providing a physical and biological barrier against pathogenic species.^49^ The tight junctions formed between densely packed epithelial cells contribute to the physical barrier, whereas the mucus and anti-bacterial peptides secreted by the cells create a biological barrier.

Measurement of transepithelial electrical resistance (TEER) is a widely adopted metric for evaluating the physical barrier of an epithelium. TEER provides a relatively non-invasive measurement of average barrier integrity by reflecting the ionic conductance of the paracellular pathway.^50^ TEER values are difficult to compare directly to *in vivo* behavior, especially since permeability is not uniform *in vivo* and local state of differentiation in a monolayer may affect average TEER values. TEER values above 300-400 Ω · cm^2^ are generally considered to reflect an intact mucosal barrier.^50,51^ To evaluate the effects of *F. prausnitzii* on the physical barrier of colon epithelium, we measured the TEER of epithelia cultured either in GuMI with no bacteria (**GuMI-NB**), in GuMI with *F. prausnitzii* (**GuMI-FP**), or in a standard incubator under static conditions (**Static**) (**Figure 2e**). We observed that as early as day 2, the static group maintained significantly higher TEER values (1412 ± 150 Ω · cm^2^) compared to GuMI-NB (482 ± 108 Ω · cm^2^, P=0.01) and GuMI-FP (616 ± 205 Ω · cm^2^, P=0.04), and a similar trend obtained at day 3. Importantly, despite the differences, the epithelial barriers remained intact in each condition with average TEER values greater than 480 Ω · cm^2^, i.e., significantly above the standard threshold reflective of intact monolayers.

The integrity of the physical barrier was further validated by inspecting the monolayers under phase-contrast microscopy and immunofluorescent confocal microscopy (**Figure 2f-i**). Colon epithelial monolayers maintained inside GuMI with constant apical flow, in both the absence and presence of *F. prausnitzii*, were intact and similar in appearance to those maintained in Static conditions (**Figure S2a-c**), containing similar distributions of differentiated cells as evidenced by staining for Muc2 (goblet cells), and NHE3 (colonocytes) (**Figure 2f-h**). The epithelium also had proper polarization of sodium-hydrogen antiporter (NHE3) on the apical surface (**Figure 2h. XZ projection**), critical for transepithelial Na^+^ absorption, intracellular pH, and nutrient absorption.^52^

In the GuMI-FP condition, a large cloud of *F. prausnitzii* cells (blue) about 100-200 um thick was observed on top of colon epithelial cells with a ∼15 µm gap area between the microbes and epithelial surface (**Figure 2i**). The gap area is likely the tightly-crosslinked inner mucus gel that prevents the bacterial cells from being direct contact with the colon cell membrane.^48^ These results, together with the muc2 staining (**Figure 2g**), suggest the bacterial cells in the GuMI device reside relatively close to the colon epithelia in an outer diffuse mucus layer, in an arrangement similar to mucus-associated microbes *in vivo*. The relatively low shear in the GuMI flow arrangement may facilitate the development of this robust mucus layer.

We next confirmed the biological barrier of the epithelium by measuring the concentration of apical mucin in GuMI-NB, GuMI-FP, and Static conditions. Although samples cultured in GuMI were constantly being perfused with anoxic apical medium, the concentration of mucin measured inside Transwells harvested at the end of the 3-day experiment was comparable between GuMI-NB (2.3±1.2 µg/ml), GuMI-FP (4.1 ± 1.6 µg/ml), and Static (3.8 ± 2.2 µg/ml) (**Figure 2j**). No mucin was detected in the effluent collected at the apical effluent reservoirs of GuMI-NB and GuMI-FP samples (**Figure 2j**), although it may have been present below the detection limit of 0.5 µg/ml. Taken together, our data suggest that GuMI addresses the metabolic needs of both obligate anaerobes and colon epithelial cells and allows epithelium to maintain its physical and biological barrier properties.

### The GuMI physiome platform supports fermentation by the super oxygen-sensitive bacterium *F. prausnitzii*

After verifying that GuMI can maintain both the colon epithelial barrier and obligate anaerobes, we next asked if bacterial fermentation occurs. *F. prausnitzii* is one of the most abundant species in the human fecal microbiota^53^ and the major producer of butyrate.^54^ It produces butyrate abundantly (>10 mM) *in vitro*.^12,55^ To determine if *F. prausnitzii* cells actively produce butyrate when co-cultured with human colon epithelia, we compared the concentration of butyrate in the apical medium and the GuMI effluent in the absence (GuMI-NB) and presence of *F. prausnitzii* (GuMI-FP). As expected, the concentration of butyrate in the apical medium in GuMI-NB (0.08±0.04 mM) or the inlet (source) medium (0.10±0.04 mM) was close to or below the detection limit 0.08 mM (**Figure 2k**). In contrast, butyrate in the bulk apical medium collected from the GuMI-FP condition increased significantly (p < 0.0001), to 0.75±0.31 mM (**Figure 2k**). We attribute this increase to the fermentation of apical medium substrates by *F. prausnitzii*. Conversely, no statistically relevant change was observed for the other two SCFA, acetate and propionate, between GuMI-NB and GuMI-FP (**Figure 2k**). A similar trend – i.e., a dramatic increase in butyrate for the GuMI-FP condition, with no or minor discernible changes in other SCFA – was observed in the GuMI-NB and GuMI-FP effluents (**Figure S2d**). These data suggest that *F. prausnitzii* is functionally active without compromising colon epithelial barrier functions (**Figure 2e-j**), as the compromise of barrier function would cause an influx of oxygen and death of the bacteria.

*In vivo*, SCFAs are transported apically from the luminal surface into colonocytes by the monocarboxylate transporter 1 (MCT1, also known as SLC16A1), where they are partially metabolized before excretion on the basolateral surface by SLC16A1 and possibly other transporters.^56^ We observed high levels of butyrate in the basolateral medium of the monolayers only from GuMI-FP, indicating that butyrate was actively taken up from the apical side and transported to the basal compartment in the presence of *F. prausnitzii* (**Figure 2l**). Under Static conditions, SCFAs were at low levels in either the basolateral or apical medium (**Figure 2k** and **2l**), suggesting that SCFAs are consumed by colon epithelium during transport as we have previously reported for microbe-free Transwell cultures of primary human colon supplemented with apical SCFAs.^57^ The relatively low concentrations in Static likely reflect a general nutrient depletion state relative to GuMI-NB and GuMI-FP, as the total volume of apical medium to which the epithelial barrier in GuMI-NB or GuMI-FP is exposed during the culture period is 28.8 mL (10 µl/min × 48 h × 60 min/h) compared to 0.5 mL over the same time period in Static culture. In the GuMI-FP condition, in contrast, the measured final concentrations of propionate and butyrate in the apical compartment were at or above the source concentration, and concentrations in the basolateral medium were unexpectedly slightly higher than those in the apical media for GuMI-FP, in contrast to the descending SCFA gradient observed from colon to peripheral blood *in vivo*.^58^ This observed phenomenon likely reflects the highly inhomogeneous distribution of microbes in the apical compartment, with microbes concentrated in the mucus region associated with the monolayer (Fig. 2i), causing SCFA fermentation products to reach locally high concentrations compared to that in the apical medium flowing above the mucus layer and driving active transport across a physiologically-normal descending gradient. The characteristic diffusion time for butyrate from the apical surface to the upper barrier in the GuMI device, a distance of 3 mm, is ∼ 2-5 hr (see Supplemental Methods A) compared to the average time of 35 min required to replace medium in the apical compartment (volume ∼350 µl) at the standard flow rate of 10 µl/min. Thus, the collection of the bulk apical medium for analysis likely dilutes the local peri-epithelial concentrations significantly.

Differences in the total volume of medium that cells experience in Static compared to GuMI culture made it difficult to directly compare the overall contribution of *F. prausnitzii* to the concentration of SCFAs over the course of the experiment using single time-point concentrations, we therefore calculated the net consumption or production (µmole) of individual SCFA over the entire course of the experiment, summarized in **Table 1**. Net consumption and transport of both acetate and propionate were observed across all conditions (i.e., Static, GuMI-NB, GuMI-FP). Net production of butyrate, however, was observed only in the presence of *F. prausnitzii* (**Table 1**). These results indicate a significant amount of butyrate was produced in apical media, while similar amounts of acetate and propionate were transported from the apical to the basolateral medium in the presence of *F. prausnitzii*. The total amounts of butyrate present in the (1) apical, (2) basolateral media, and (3) apical effluent at the end of the experiment in GuMI-FP were 0.26, 1.6, and 22.7 µmole, respectively, representing 5-8 times higher amounts compared to these three compartments in the GuMI-NB.

**Table 1.** Net consumption or production (µmole) of SCFA acetate, propionate, and butyrate in the apical medium, basolateral medium, and apical effluent under Static, GuMI-NB, and GuMI-FP conditions.

| SCFA | Apical medium |  |  | Basolateral medium |  |  | Apical effluent |  |  |
| --- | --- | --- | --- | --- | --- | --- | --- | --- | --- |
|  | Static | GuMI-NB | GuMI-FP | Static | GuMI-NB | GuMI-FP | Static | GuMI-NB | GuMI-FP |
| Acetate | -0.8±0.02a | -0.2±0.1a | -0.2±0.08a | 0.3±0.4a | 1.5±0.6b | 1.4±0.6b | NA | -24±45a | -21±38a |
| Propionate | -0.3±0.03a | -0.05±0.06a | -0.08±0.07a | 0.3±0.3a | 2.0±0.7b | 2.1±0.5b | NA | -13±15a | -4.6±16a |
| Butyrate | -0.02±0.01a | -0.001±0.03a | 0.2±0.1b | 0.07±0.08a | 0.2±0.1b | 1.6±0.3c | NA | 0.1±3a | 19±10b |
| Total | -1.1±0.05a | -0.2±0.1b | -0.08±0.1c | 0.7±0.6a | 3.5±0.8b | 5±0.6c | NA | -37±48a | -6±60a |

In the human large intestine, the molar ratio of acetate, propionate, and butyrate is around 3:1:1.^59^ A mixture of these SCFA, especially propionate and butyrate, may have additive effects on their biological activities in human colon cells.^60^ While the total cumulative concentrations of SCFA in the apical GuMI-FP compartment (4 mM) were lower than *in vivo* (90 mM),^61^ *F. prausnitzii* adjusted the molar ratio of acetate, propionate, and butyrate from its original value in the source, 2.4:1:0.15 (**Figure 2k, Table S2**), to the 3:1:1 ratio observed *in vivo* in human large intestine,^58^ suggesting that GuMI-FP can faithfully capture some features of SCFA exposure in human colon. Finally, we note that the production of butyrate is not likely limited by the diffusion rate of glucose into the microbial layer associated with the mucus (see Supplemental Method B).

### Overview of the impact on gene expression induced by GuMI and bacteria

Physiological cues such as oxygen gradients, luminal flow, commensal bacteria, and SCFA created in the GuMI physiome platform contribute to the phenotype of the epithelial barrier. To further understand the molecular effects of these microenvironmental cues on human colon epithelia, we performed mRNA sequencing on the cells harvested from monolayers under three conditions: Static, GuMI-NB, and GuMI-FP.

To get an overview of the effects of both GuMI physiome platform and *F. prausnitzii*, differential gene expression analysis was performed for the comparison of two groups: GuMI-NB *vs*. Static, and GuMI-FP *vs*. GuMI-NB (**Figure S3**). More than 24000 genes were included for both comparisons, with the exclusion of zero-expression genes (Transcripts Per kilobase Million = 0). Compared to the Static, the expression levels of 1627 genes were significantly changed (adj. p<0.05, |log_2_Fold Change| >0.5. **Figure S3a** and **Table S3**) in GuMI-NB, accounting for 6.7% of the expressed genes. Among the genes changed in the GuMI-NB condition, 796 genes were increased, while 831 genes were decreased (**Figure S3c**).

Unexpectedly, 4834 genes were significantly higher in cells in GuMI-FP relative to GuMI-NB (**Figure S3b** and **Table S4**), accounting for 20% of the expressed genes. Of these 4834 genes, more than 4000 were changed solely by the presence of *F. prausnitzii* (**Figure S3c**), as they were unchanged between GuMI-NB and Static. Only a small fraction (<16%) of the *F. prausnitzii*-influenced genes overlapped with GuMI-influenced genes (**Figure S3c**), indicating that the effects of culture alone on colon epithelia in the GuMI platform are very different from that of co-culture with *F. prausnitzii* at the transcriptional level. The altered genes are broadly distributed to all chromosomes, suggesting an extensive and non-chromosome-specific influence on gene expression (**Figure S3d** and **S3e**). While the differences between GuMI-NB and Static are expected due to differences in oxygen gradients across the monolayer, apical nutritional exposure, and effects of fluid flow other than nutrition (low-level shear stress, and washout of autocrine factors produced at the apical surface), such differences were more profound than expected. Comparative changes in gene expression in germ-free animal models *in vivo* compared to those colonized with *F. prausnitzii* are not available, as efforts to colonize germ-free animals with *F. prausnitzii* as monocultures have been unsuccessful and colonization in animals hosting *B. thetaiotaomicron* requires multiple introductions over several weeks.^17^

### GuMI accelerates cell differentiation and represses proliferation with no evidence of apoptosis

Next, we asked what molecular pathways in the colon epithelia are influenced by these physiologically relevant stimuli, i.e., apical hypoxia and continuous flow of the fresh apical medium, in the absence of bacteria, as the fluid flow is associated with enhanced differentiation of some stem cells.^62^ However, SCFAs, particularly butyrate, also suppress cell proliferation in colon epithelial cells and the greater exposure to butyrate in GuMI culture via continuous apical medium replenishment of butyrate-containing medium may also alter differentiation and proliferation.^63^ Differential gene expression analysis indicates a significant upregulation of representative cell differentiation marker genes, e.g., SLC26A3, KLF4, CDX1, DLL1, CEACAM7, and CEACAM6 (**Figure 3a**). Consistently, most of these genes, SLC26A3, CDX1, DLL1, and CEACAM7 were found to be increased in more differentiated colon monolayers derived from other human donors.^33^ Meanwhile, proliferative or stem cell marker genes such as MKI67, LGR5, ASCL2, and MYBL2 were significantly decreased (**Figure 3a**). RT-qPCR confirmed a subset of these proliferative markers, MKI67 and LGR5, were decreased, while differentiation markers SLC26A3, CDX1, and CEACAM6 were increased (**Figure 3b**). These results suggest that cells in the GuMI are more differentiated than cells cultured in Static condition. To further support this idea, gene set enrichment analysis (GSEA) was performed. GSEA is designed to detect coordinate changes in the expression of genes that are related in terms of biological function, chromosomal location, or regulation.^64,65^ This approach has been widely used since its introduction in 2003.^65,66^ Herein, GSEA takes into consideration the genes that are known to be related to cell differentiation and therefore gives a comprehensive evaluation of the cell differentiation status. GSEA revealed that the cell differentiation gene set is over-represented in GuMI-NB cells, with more than 150 genes being enriched in GuMI-NB (**Figure 3c** and **Table S5**).

**Figure 3.**
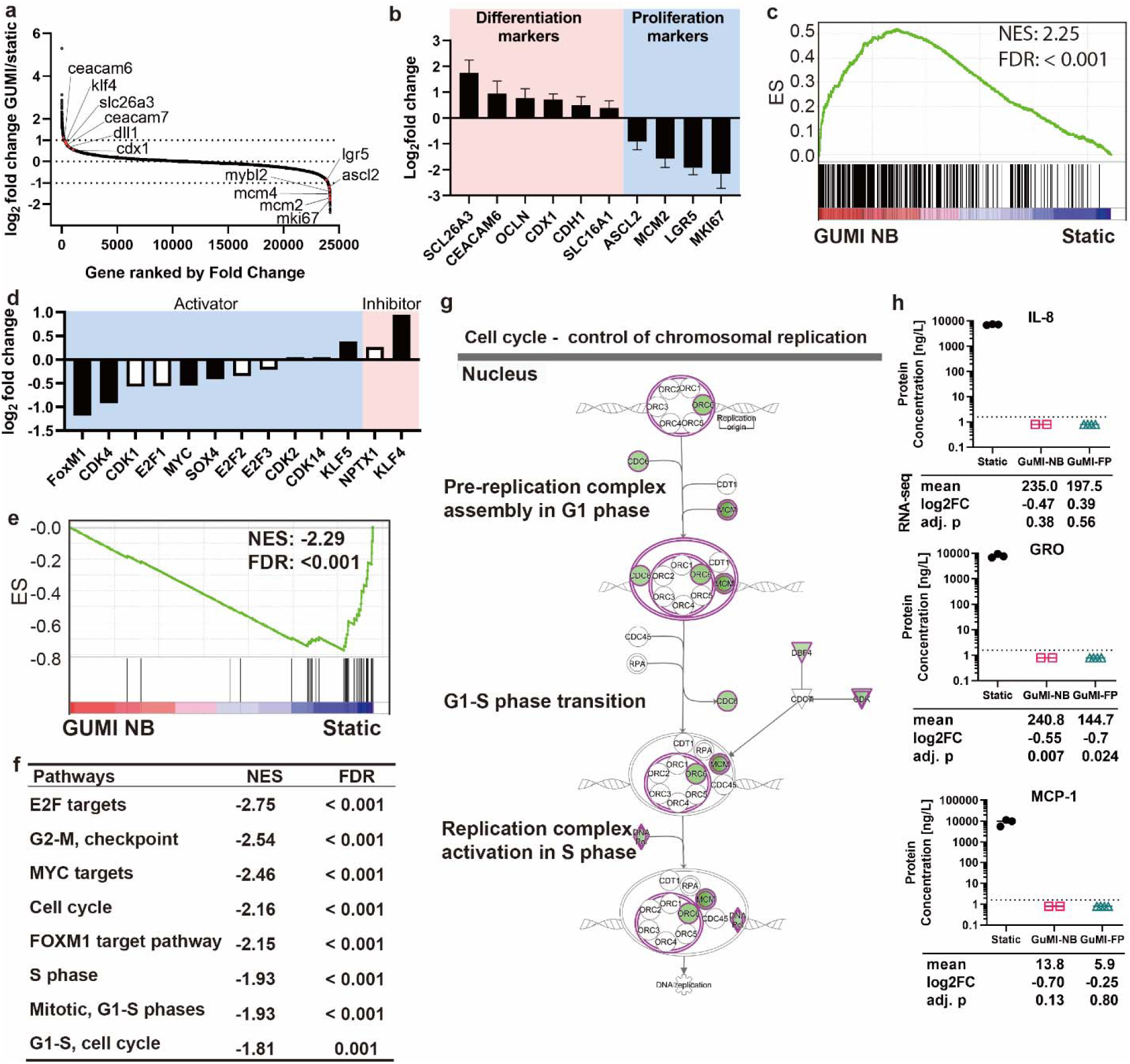
GuMI promotes cell differentiation and represses cell proliferation. (**a**) Increased expression of representative differentiation marker genes (left upper edge) and decreased expression of representative stem cell or cell proliferation marker genes (bottom right edge) in GuMI vs Static cells. All marker genes are highlighted in red dots; all the gene symbols were in small letters due to space limit. (**b**) RT-qPCR confirmation of cell differentiation and cell proliferation marker genes, bars represent mean values from 6-8 replicate samples from 3-4 independent experiments. (**c**) Gene set enrichment analysis (GSEA) for the cell differentiation gene set revealed an over-representation in GuMI over Static monolayers. NES: normalized enrichment score. FDR: false discovery rate. (**d**) Fold change in the expression of cell proliferation regulatory genes in GuMI over Static monolayers. Black-filled bar indicates a significant increase or decrease; white bar indicates a non-significant increase or decrease. (**e**) GSEA for DNA synthesis (cell proliferation) is repressed in GuMI cells. NES: normalized enrichment score. FDR: false discovery rate. (**f**) Pathways related to the control of cell cycle and targeted by transcriptional factors MYC, FOXM1, and E2F were under-represented in GuMI cells, analyzed by GSEA. (**g**) Ingenuity Pathway Analysis (QIAGEN Bioinformatics) revealed the repression of DNA replication processes. Green background color indicates a significant downregulation, and white refers to no significant change. (**h**) Concentration of IL-8, GRO, and MCP-1 at protein levels in apical media from Static, GuMI-NB, and GuMI-FP.

In the normal healthy colon, the more differentiated cells in the lumen are non-proliferative relative to the stem and progenitor compartment in the crypt, suggesting that GuMI-NB culture conditions may suppress proliferation. Cell proliferation is regulated by multiple transcriptional factors and kinases.^67^ Some transcriptional factors such as MYC, FOXM1,^68^ MYBL2, SOX4, SOX9,^69^ CDK4,^70^ and KLF5 act as activators of cell proliferation, while others such as KLF4 and LGALS1^71^ are inhibitors of cell proliferation. Indeed, the proliferation marker genes were significantly reduced in GuMI-NB culture (**Figure 3b**), suggesting that the promotion of cell differentiation was accompanied by the inhibition of cell growth and proliferation. We observed a significant reduction of activators such as FOXM1 and CDX1, and an increase of cell proliferation inhibitors such as KLF4 (**Figure 3d**). For example, FOXM1 stimulates proliferation by promoting S-phase entry as well as M-phase entry and is involved in the proper execution of mitosis.^72^ These processes involve DNA synthesis, which is a critical preparation process for cell proliferation. Therefore, we performed GSEA for the genes in DNA synthesis machinery. The results clearly indicated that the DNA synthesis pathway is over-represented in Static cells and dramatically repressed in GuMI-NB cells (**Figure 3e**). The core genes for the DNA synthesis pathway were significantly decreased in GuMI-NB cells. These genes are responsible for the pre-initiation and initiation of DNA synthesis, elongation and maturation of newly synthesized DNA (**Figure S3f**). In addition, other pathways related to cell cycle regulation and MYC/FOXM1/E2F-target pathways were found to be significantly repressed in GuMI-NB cells (**Figure 3f**). Ingenuity Pathway Analysis (IPA) confirmed that the processes for controlling chromosomal replication in the nucleus were decreased in GuMI-NB (**Figure 3g**). This is potentially due to the downregulation of proliferation-associated transcription factors. Indeed, using upstream regulator analysis in IPA, an algorithm that identifies molecules upstream of a gene set or pathway, we identified three proliferation-activating transcription factors, MYBL2, FOXM1, and MYC (**Table S6**) and proliferation-inhibitory factors LGALS1 and KLF4 (**Table S6**). In addition, the WNT-activating gene SOX4^73^ is decreased in GuMI cells. SOX family actively interacts with the WNT/β-catenin pathway, which is key to maintaining the colonic stem cells.^74^

We did not see enhancement of apoptotic genes (e.g., apoptosis-inducing factor AIFM1, AIFM2, AIFM3, DIABLO [also known as SMAC], apoptosis regulator BCL2) nor did we observe evidence of death or apoptosis in the monolayers via microscopy. Together, these results suggest that the physiological stimuli maintained by the GuMI physiome platform promotes cell differentiation and represses cell proliferation, likely owing to the changes in multiple transcription and growth factors.

### GuMI-NB modulates apical autocrine factors associated with proliferation, differentiation, and inflammatory responses, and co-culture with *F. prausnitzii* is associated with further changes

The transcriptional changes in differentiation and proliferation genes between Static and GuMI-NB culture imply changes in proteins regulating these processes. Epithelial mucosal barriers regulate their homeostasis and response to injury in part by polarized production of numerous growth factors, chemokines and cytokines.^75^ Most of these factors act not only in paracrine fashion to recruit immune cells after injury and signal neighboring stromal cells, but also in autocrine fashion: colon epithelial cells express not only canonical growth factor receptors (e.g., epidermal growth factor receptor, EGFR; fibroblast growth factor receptors, FGFRs; platelet-derived growth factor receptors, PDGFRs) but also receptors for chemokines (CXCR1-4; CCR2-5) and cytokines (receptors IL-1, IL-4, IL-15, and IL-18).^76-86^ Autocrine loops regulate colon epithelial barrier permeability, proliferation, response to infection, and diverse other behaviors, and in turn, the activity of autocrine loops is influenced by gut microbes.^75^ Compared to static culture, where autocrine factors accumulate over 48 hr between medium changes, continuous flow in the apical compartment of GuMI, where the 0.35 mL medium volume is replaced about twice an hour, dilution of autocrine factors may influence autocrine signaling loops and the resulting phenotype. The presence of *F. prausnitzii* may further influence autocrine loop activity.

We therefore compared concentrations of 47 growth factors, cytokines, and chemokines in the medium collected from the apical compartment of Static, GuMI-NB and GuMI-FP after two days of co-culture, using multiplex immunobead assays to assess protein concentrations. The mucus layer was included in the collected samples. Twenty five of the 47 proteins probed were detectable above background levels in Static cultures (**Figure 3h** and **Figure S4**). Chemokines IL-8, GRO (CXCL1), and MCP-1 (CCL2), which are known to be produced at high levels on the apical surface to signal breaches of the epithelial barrier for neutrophil recruitment and are also autocrine factors, were observed at 600 - 900 pM (6,000 – 10,000 pg/mL) -- more than an order of magnitude greater than other detected proteins (**Figure 3h** and **Figure S4a-d**). Ten factors with autocrine signaling properties in colon epithelia were detected at concentrations of 3 – 20 pM, including (in order of descending concentration) TNF-α, RANTES, GM-CSF, IL-1RA, PDGF ab/bb, IL-4, TGF-α, FGF-2, G-CSF, and fractalkine (**Figure S4a**). Twelve additional factors, including IL-15, IL-10, IL-13, IL-1α, IL-7, IL-21, IL-18, IP-10, Eotaxin, IFN-α2, MIP-1α, and sCD40L (**Figure S4b**), accumulated to concentrations in the 0.3-2 pM range, whereas 20 factors were undetectable above background (**Figure S4c-d**). These values are comparable to others reported for IL-8 and other overlapping cytokines in *in vitro* colon epithelia cultures.^75^ Chemokines, cytokines, and growth factors exert signaling activities at concentration thresholds in the 5-100 pM range. While concentrations of some of these factors as measured in the bulk apical media are borderline or below those expected to cellular responses, they may be sufficient to act in autocrine fashion as concentrations of autocrine factors measured in the bulk are known to underestimate the local concentrations at the cell surface, sometimes dramatically so.^87^

None of the 47 factors were robustly detected above background levels in the apical media collected from either GuMI-NB or GuMI-FP at the same 48 hr time point (**Figure 3h, Figure S4a-d**), which is not surprising considering the apical media is diluted more than 100-fold compared to the static culture over 48 hr. However, autocrine factors are likely strongly concentrated at the cell surface in the GuMI device as a 100-200 µm quiescent mucus layer presents a diffusion barrier between the cell surface and the bulk apical flow. For example, assuming comparable production rates between Static and GuMI-NB, we estimated that IL-8 and TGF-α would reach cell surface concentrations of ∼7 pM and 0.1 pM, respectively (see **Supplemental Method C**). Indeed, it is unlikely that cellular production rates of IL-8, MCP-1, and most other factors changed dramatically between flow and static conditions, as only one of the 27 factors detected at the protein level, GRO (CXCL1), showed significant changes at the transcriptional level between GuMI-NB and Static, exhibiting downregulation under flow conditions (**Figure S4a-d**). CXCL1 is upregulated in inflammation and colon cancer, and its receptors CXCR1 and CXCR2 are both expressed in vivo on colon epithelia,^76,80^ suggesting a role in proliferation, but the direct effects of CXCL1 or other CXCR1 and CXCR2 ligands (e.g. IL-8) on primary human colon epithelial cell proliferation have not been investigated to our knowledge, and receptor expression in cell lines does not appear to reflect that observed *in vivo*.^80^ Activation of CXCR1 and CXCR2 in several epithelial cancer cell types initiates multiple downstream signaling pathways including MAP (mitogen-activated protein) kinase and phosphoinositide 3-kinase which stimulation migration and proliferation.^88^ Two other factors, IL-1β and IL-33 (**Figure S4d**), which were not detected at the protein level, showed significant upregulation and downregulation, respectively, in GuMI-NB compared to Static, while 6 other factors not detected at the protein level in GuMI-NB showed no change at the transcriptional level between GuMI-NB and Static conditions. IL-33 is an alarmin stored in the nucleus which is upregulated by EGFR activation,^89^ suggesting a lower activation state of EGFR consistent with the reduced concentration of the EGFR ligand TGF-α in GuMI-NB compared to Static. It is unclear if the regulation of IL-1β, a cytokine upregulated in inflammation, plays a major functional role as its antagonist, IL1-RA, is robustly expressed (**Figure S4a**). Overall, the apparent reduction in stimulatory cytokines at the apical surface is consistent with the reduction in proliferative signatures.

The presence of *F. prausnitzii* in the GuMI-FP condition induced transcriptional changes in 6 proteins compared to the GuMI-NB condition: GRO and IL-33, which were both downregulated in GuMI-NB compared to Static, were further downregulated in GuMI-FP; FGF-2 was downregulated and TNF-α, IL-15, and IL-18 were upregulated in GuMI-FP compared to GuMI-NB. Downregulation of FGF-2, which stimulates proliferation in colon stem cells,^82^ is consistent with the much greater exposure of cells in GuMI-FP compared to those in GuMI-NB to butyrate (Fig. 2 k-l), an inhibitor of intestinal cell proliferation. Upregulation of TNF-a is difficult to interpret, as it acts in a highly context-dependent manner, enhancing cell death and inflammation in some contexts and stimulating cell survival and proliferation in others.^85,90^. Similarly, direct effects of IL-15 on colon epithelia are unclear; its production is associated with paracrine regulation of cell survival,^91^ and the Caco-2 cell line shows activation of downstream signaling by IL-15^92^ but human intestinal epithelia do not express functional IL-15 receptors, only splice variants unable to bind IL-15.^93^ Upregulation of IL-18, a cytokine essential for gut barrier integrity,^94^ is consistent with reported effects of butyrate on gut barrier function.^95^ Overall, the pattern of changes between GuMI-FP and relative to GuMI-FP suggest improved homeostatic functions consistent with enhanced exposure to butyrate.

### GuMI recapitulates cell responses to hypoxia

Colon epithelial cells are exposed to a uniquely steep oxygen gradient. This oxygen gradient helps maintain a healthy colon microbiota comprising many oxygen-sensitive commensals,^96^ preventing aerobic pathogen expansion,^97^ and maintaining host homeostasis.^96^ Hypoxia-induced factor-1 alpha (*HIF1A*) is a master transcriptional regulator of cellular response to hypoxia.^98,99^ Thus, we hypothesized that apical hypoxia in GuMI culture would induce hypoxia-related responses in colon epithelia and promote epithelial barrier functions. First, we found under hypoxic conditions *HIF1A* gene expression was significantly increased (p = 0.0001) in GuMI-NB compared with Static cells cultured in normoxia, with a fold change of 1.52 (**Figure 4a** and 4**b**). RT-qPCR validation confirmed the fold change is about 1.80 (**Figure 4c**). The stability of HIF1A protein is regulated by an ubiquitin-proteasome-based degradation, which mandatorily requires oxygen and HIF1A hydroxylases (EGLN1-3 and HIF1AN).^100^ The transcription of HIF hydroxylases EGLN1-3 and HIF1AN was not significantly changed in GuMI-NB (p > 0.4 for all genes in **Supplementary Figure S4e**), together with the hypoxic conditions maintained in GuMI-NB (**Figure 1b**), suggested that HIF1A is likely stabilized.

**Figure 4.**
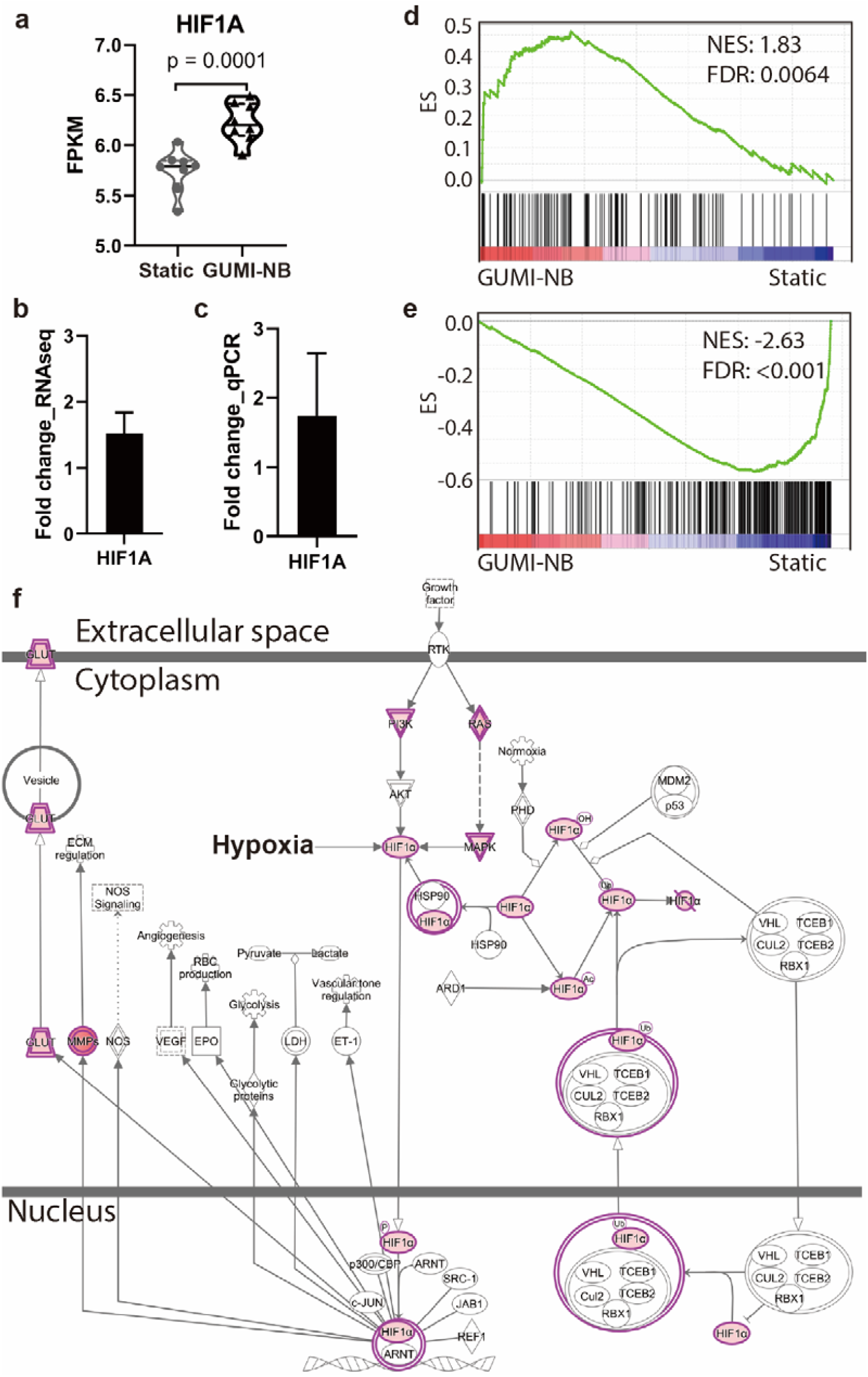
The physiological oxygen gradient increases hypoxia sensing pathway. (**a**) increased expression of *HIF1A*, a hypoxia response gene. Fold change of *HIF1A* mRNA based on (**b**) RNA-seq and (**c**) RT-qPCR. Bars in (**b-c**) represent the mean±SD of 6-8 replicate samples. (**d**) GSEA revealing that genes in response to HIF1A are overrepresented in GuMI cells. (**e**) GSEA revealing that genes downregulated in response to HIF1A are underrepresented in GuMI cells. (**f**) IPA revealing the curated pathways responding to HIF1A are increased in response to increased expression of *HIF1A*.

HIF1A is a heterodimeric DNA-binding complex that regulates an extensive transcriptional response to hypoxia.^101^ The activation of HIF1A induced by hypoxia has a protective role in mucosal barrier function *in vivo* and on cells *in vitro*.^102^ We thus asked if the target genes or pathways of HIF1A are changed due to elevated expression of HIF1A. To test this, we performed GSEA using the compiled gene sets that are known to respond to HIF1A or HIF1A inducers in different types of cells.^103,104^ Genes known to be increased in response to HIF1A are indeed over-represented in GuMI-NB, while genes known to be decreased by HIF1A are underrepresented in GuMI-NB (**Figure 4d** and **4e**). This response agrees with Ingenuity Pathway Analysis, which revealed that pathways responding to HIF1A are significantly upregulated (**Figure 4f**). Together, these results suggest that hypoxia conditions maintained by GuMI are able to recapitulate colonic cellular responses to a physiologically-steep oxygen gradient. The increase of *HIF1A* mRNA, and suppression of cell cycle related pathways was also observed in Caco-2 monolayers cultured anaerobically with *F. prausnitzii* for 12 h.^23^

### *F. prausnitzii* exerts anti-inflammatory effects on the epithelial monolayer and represses TLR3 and TLR4 expression

The GuMI device maintains a microenvironment that supports chronic (2-4 days) co-culture of a primary human colon monolayer with a continuously-growing commensal bacterial population including the strictest of anaerobes *F. prausnitzii* (**Figure 2**). *F. prausnitzii* is one the most abundant species in the human colon microbiome. In humans, *F. prausnitzii* is the major producer of butyrate (**Figure 2k**) which, together with other metabolites,^14^ mediates an array of effects on host cells including modulation of immune cell behavior and inhibition of proliferation and inflammation.^96,105-107^ Secreted products of *F. prausnitzii* have been shown to modulate the NFKB signaling pathway in both cell culture and mouse colon injury models.^9,46^ *F. prausnitzii* has been identified as an anti-inflammatory bacterium, whose abundance was significantly decreased in IBD patients.^9,108^ To further test if GuMI-FP was able to recapitulate the anti-inflammatory effects of *F. prausnitzii*, we examined the transcription of NFKB1, its upstream regulation pathway and its downstream target genes. NFKB1, the key subunit of the NFKB complex, was significantly decreased by two-fold in GuMI-FP over GuMI-NB cells. The NFKB complex is regulated by multiple mechanisms, including NFKB inhibitors, NFKB activators, and TLR-NFKB signaling. We first examined the gene expression of NFKB inhibitor (NFKBI) genes, which encode NFKBI proteins to form a complex with RELA and NFKB1, preventing the activation of NFKB. To activate NFKB, IkappaB kinase (IKBK) phosphorylates NFKI protein, which then is ubiquitinated and degraded by proteasome.^109^ Herein we show that four out of five NFKBI genes, i.e. NFKBIZ, NFKBIA, NFKBIE, and NFKBIB, were increased (**Figure 5a** and **5b**). On the other hand, IKBKE, one of the three NFKB activating genes, was decreased, while the other two were not significantly changed (**Figure 5a** and **5b**).

**Figure 5.**
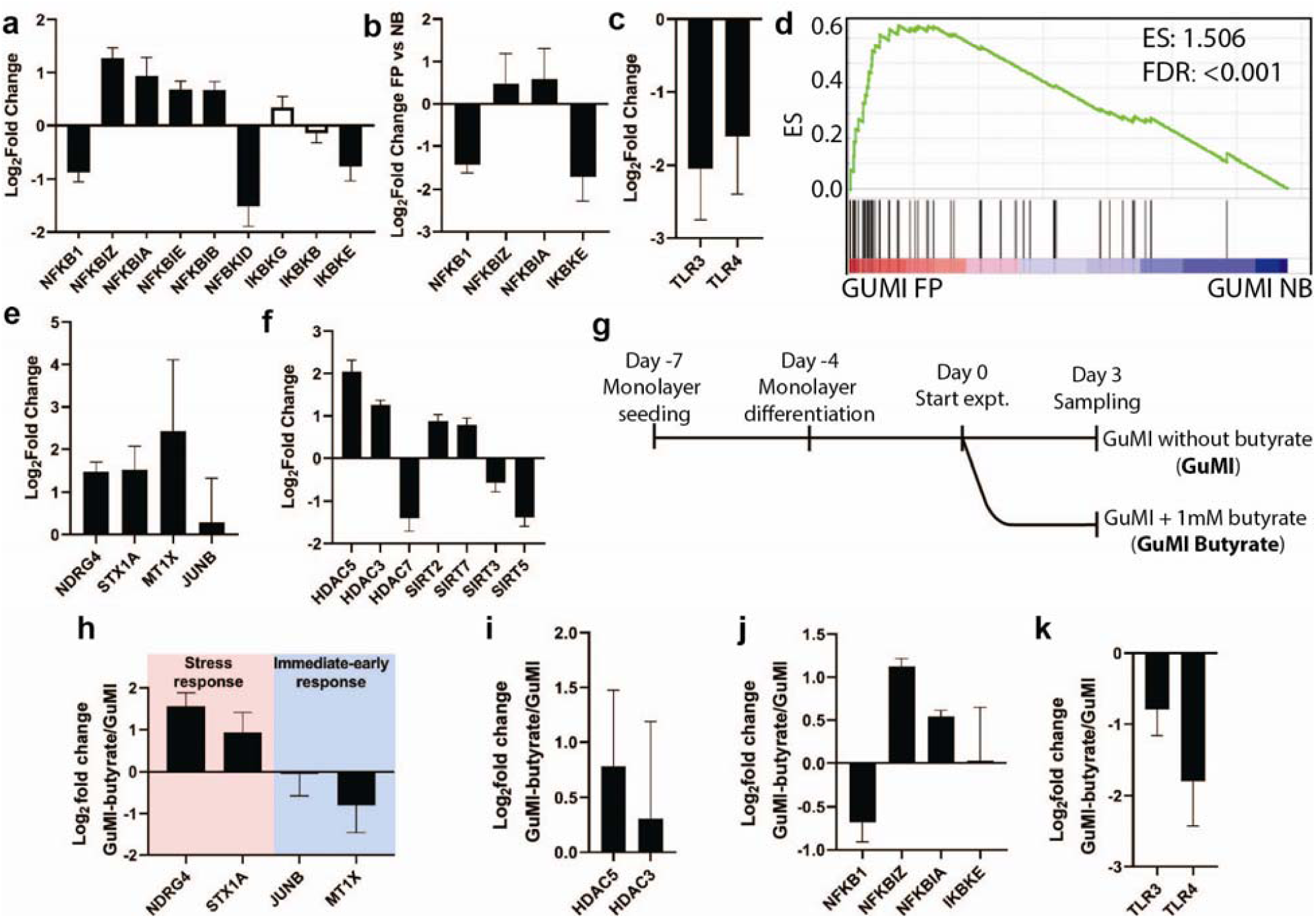
Effects of *F. prausnitzii* on colon epithelia in GuMI. (**a**) RNA-seq analysis reveals changes in gene expression of NFKB1, its inhibitors (NFKBI) and activators (IKBK) induced by *F. prausnitzii*. (**b**) Validation by RT-qPCR of changes in NFKB1, NFKBIZ, NFKBIA, and IKBKE induced by *F. prausnitzii*. (**c**) Validation by RT-qPCR of changes in TLR3, TLR4 induced by *F. prausnitzii*. (**d**) GSEA revealing that genes that respond to butyrate are overrepresented in GuMI-FP cells over GuMI-NB cells. Differentiation and proliferation gene sets are not overrepresented. (**e**) Validation by RT-qPCR of a subset of butyrate responsive genes. (**f**) RNA-seq analysis reveals HDAC genes that are significantly (log2FC ≥ 0.5, adj. p <0.05) changed by *F. prausnitzii*. Bars in (**a-c, e-f**) represent the Mean±SD of 4-8 replicate samples from 3-4 independent experiments. (**g**) Timeline of the experiment to determine the effects of butyrate on colon epithelial cells in GuMI. (**h-k**) RT-qPCR analysis of (**h**) a subset of butyrate-responding genes NDRG4, STX1A, JUNB, and MT1X; (**i**) HDAC3 and HDAC5; (**j**) NFKB1, NFKBIZ, NFKBIA, and IKBKE; (**k**) TLC3 and TLR4 in GuMI-butyrate over GuMI. Bars represent the Mean±SD of 3 replicate samples from 2 independent experiments.

We next looked at the toll-like receptors (TLR), including TLR2, TLR3, and TLR4, which are directly involved in the molecular recognition of bacteria and regulation of the NFKB pathway.^110^ and which show dysregulated expression in UC and CD patients.^18^ We find that *F. prausnitzii* down-regulates TLR3 and TLR4 expression and the NFKB pathway in primary colon epithelia (**Figure 5c)**. This finding of reduction in TLR4 expression is consistent with *F. prausnitzii* as more abundant in healthy individuals than patients with UC or Crohn’s;^9,108^ however, TLR 3 is reduced in active CD compared to healthy, and not changed in UC patients.^18^ Using Caco-2 cells, it was found that *F. prausnitzii* activates TLR3 in Caco-2 cells during their short-term and aerobic co-culture.^111^ Further studies are needed to illuminate the discrepancies in TLR3.

Two pathways can respond to changes in TLR3 and TLR4. One is dependent on MYD88, TIRAP, and IRAK, and the other one is dependent on TICAM and MAP3K14.^112,113^ These two responding pathways upregulate the same downstream factors, i.e. MAP3K7 and TAB, which then regulate the phosphorylation of NFKB. Interestingly, with the decrease of TLR3 and TLR4 (**Figure 5c)**, the responding pathway MYD88-TIRAP-IRAK was downregulated, whereas the other responding pathway TICAM1-MAP3K14 was up-regulated. As a result, MAP3K7-TAB was not significantly changed (**Figure S4f**). Together, these results suggest that *F. prausnitzii* inhibits NFKB, potentially through upregulation of NFKB inhibitors, downregulation of NFKB activator and TLR3/4, but with no change in expression of MAP3K7-TAB.

In our experimental setting, the genes known to be induced by butyrate^114-116^ are overrepresented in GuMI-FP over GuMI-NB cells (**Figure 5d**), with a subset of genes, i.e., NDRG4, STX1A, and MT1X, in the gene set confirmed by RT-qPCR (**Figure 5e**). Consistently, IPA identified butyric acid as an upstream regulator which activates several pathways, including histone deacetylases (HDAC) (**Table S7**). The transcription of genes is largely regulated by epigenetic modification mediated by histone acetylases (HAT) and HDAC. Therefore, we examined the changes of HAT and HDAC genes influenced by *F. prausnitzii*. Interestingly, no HAT was changed in the presence of *F. prausnitzii*, whereas seven out of eighteen HDAC (HDAC1-11, SIRT1-7) were significantly changed (adj. p <0.05, log2FC ≥ 0.5, **Figure 5f**). HDAC3, HDAC5, SIRT2 and SIRT7 were increased, while HDAC7, SIRT3 and SIRT5 were decreased (**Figure 5f**), indicating global and differential changes across different HDAC by *F. prausnitzii*.

Despite the relatively greater exposure to butyrate produced by *F. prausnitzii*, cell differentiation was not further enhanced in GuMI-FP over GuMI-NB, as most of the differentiation marker genes did not change significantly (adj. p >0.05) in GuMI FP over GuMI-NB. These genes include SLC26A3, KLF4, CDX1, DLL1, CEACAM7, and CEACAM6. Consistently, proliferative markers were also relatively unchanged between GuMI-NB and GuMI-FP, i.e. MKI67, LGR5, and ASCL2, with one exception, MYBL2, which is significantly increased in GuMI-FP. These results indicate that the monolayer in the platform was at a very high differentiation state.

### Butyrate contributes to the anti-inflammatory effect induced by *F. prausnitzii*

To further understand whether butyrate contributes to the effects mediated by *F. prausnitzii*, we exposed cells in GuMI-NB to 1 mM butyrate in the anaerobic apical medium (**Figure 5g**). The concentration used was to mimic the production of *F. prausnitzii* in GuMI-FP. (**Figure 2k**). We first determined the expression of multiple stress response genes (NDGR4, STX1A, JUNB, and MT1X), which appears in enriched butyrate-responding pathway (**Figure 5h**), and found NDRG4 and MT1X were increased by up to three-fold, while the immediate-early response gene JUNB was not changed and MT1X was decreased (**Figure 5h**). This confirmed that butyrate contributes to the effects produced by *F. prausnitzii*, but the pattern is not exactly the same as for *F. prausnitzii*. Similar discrepancies were observed for HDAC, with a significant increase of HDAC5 and no change of HDAC3 in GuMI-Butyrate comparing to GuMI (**Figure 5i**). Interestingly, we found NFKB pathway was modulated by butyrate in a similar pattern as by F. prausnitzii (**Figure 5j** and **5a**). Finally, TLR3 and TLR4 were decreased by butyrate (**Figure 5k**) in a similar manner to the exposure to *F. prausnitzii*, suggesting that butyrate largely contributes to the TLR and NFKB regulation by *F. prausnitzii*. These results together confirmed that butyrate contributes to the gene-regulatory effects, especially TLR downregulation, induced by *F. prausnitzii*.

Butyrate (1 mM) decreases TLR4 expression in the colon cancer cell line HCT116 at 48-72 hours post-exposure.^117^ This inhibitory effect of butyrate on TLR4 expression was also observed in mouse adipose tissue *in vivo*.^118^ Sodium butyrate significantly decreased the TLR4 and NFKB signaling pathways in lipopolysaccharide-induced acute lung injury in mice.^119^ In addition, butyrate inhibits the NF-kB pathway in the HT-29 colon cancer cell line.^120^ However, this effect might be tissue- or cell type-specific. In SW480 cells and mouse colon cancer CT26 cells, butyrate upregulates TLR4 expression.^121^

The similarity in the transcriptional changes that occurred in response to butyrate and *F. prausnitzii* indicates that butyrate is an important contributor to the effects of *F. prausnitzii*. However, there are also discrepancies observed between butyrate and *F. prausnitzii*, suggesting that there are other effectors (metabolites, secreted proteins, or bacterial cell components) that contribute to the effects of *F. prausnitzii*. Indeed, *F. prausnitzii* can secrete the anti-inflammatory protein MAM,^46^ which downregulates the NFKB pathway when it is transfected into a colon cancer cell line.^46^ In another study using a gnotobiotic mouse colitis model, many metabolites in the gastrointestinal tract were associated with colitis-protective effects by *F prausnitzii*.^14^ *F prausnitzii*-produced salicylic acid, in addition to butyrate, was shown to contribute to the attenuation of inflammation.^14^

Butyrate is also a known modulator of HDAC genes in colon cancer cell lines.^106,116,122^ It has been shown that butyrate regulates the SIRT family of deacetylases in human neuronal cells, with increased expression of SIRT1, SIRT5, and SIRT6 and downregulation of SIRT2, SIRT4 and SIRT7.^123^ The regulation pattern of SIRT, the class III HDAC, is consistent with that of butyrate.^123^ However, the activation of HDAC3 and HDAC5 by *F. prausnitzii* and/or butyrate does not agree with previous observations, where both HDAC3 and HDAC5 were inhibited by butyrate.^124^ While the detailed mechanisms require further research, it is worth noting that most of these studies were carried out with cancer cell lines.^124^ Indeed, even with different colon cancer cell lines, the response to the same amount of butyrate is different, with some cell lines being very sensitive to butyrate, and others resistant.^115^ The regulation of HDACs by butyrate in other cell types is similarly heterogeneous: 1 mM of butyrate was shown to induce the expression of HDAC1 and HDAC3 in human PBMCs after 6 and 48 h, but repress the expression of HDAC2 after 48 h;^125^ oral butyrate for 8 weeks decreased HDAC2 expression in cardiac cells from Wistar rats;^126^ and sub-mM butyrate inhibits the NFKB signaling pathway and histone deacetylation in intestinal epithelial cells and macrophages.^127^

## Conclusions

This report is, to the best of our knowledge, the first to describe chronic co-culture of the super oxygen-sensitive anaerobe *F. prausnitzii* with a primary human colon mucosal barrier using continuous flow in the apical compartment to foster robust growth and metabolic activity of the microbial population. To accomplish this, we designed and fabricated a GuMI physiome platform that maintains a mucosal barrier-anaerobe interaction in a manner that allows for continuous media exchange in both compartments, along with the introduction of microbes, including pathogens, following stabilization of the epithelial barrier. This platform houses six independent culture chambers with associated independent media flow circuits. Using primary human colon epithelial monolayers co-cultured continuously with super oxygen-sensitive bacterium *F. prausnitzii* for two days, we demonstrated the epithelium experiences a steep oxygen gradient with a complete anaerobic apical environment fostering the growth and active fermentation of *F. prausnitzii*. Using transcriptomics, GSEA, and RT-qPCR, we identified elevated differentiation and hypoxia-responding genes and pathways in the platform over conventional static culture. We further used this platform to elucidate the responses of primary colon epithelia to commensal *F. prausnitzii* and demonstrated an anti-inflammatory effect of *F. prausnitzii* through the HDAC, TLR-NFKB axis. Finally, we identified butyrate largely contributes to these anti-inflammatory effects by downregulating TLR3 and TLR4. Although monolayers were used for this initial demonstration, the device can also accommodate 3D crypt-containing tissue-engineered mucosal barrier structures such as those previously described.^128^ Overall, this platform faithfully recapitulates several important features of the physiological microenvironment in the colon *in vivo* and has the potential to enable a better understanding of human colon mucosal barrier-bacteria interactions. Given the reported difficulties on translating microbiome/bacteria-based therapy from animal models to human recently,^129,130^ GuMI could help better understanding the inconsistency on the translation.

## Materials and Methods

### Chemicals and Reagents

3-nitrophenylhydrazine (N02325G, Fisher Scientific), *N*-(3-dimethylaminopropyl)-*N*′-ethylcarbodiimide HCl (50-848-678, Fisher Scientific), butyric acid (B103500-100ML, Sigma-Aldrich), propionic acid (P1386, Sigma-Aldrich), and acetic acid (338826, Sigma-Aldrich), butyric acid-d_7_ (D-0171, CDN isotope), acetonitrile (34998, Sigma-Aldrich). Ultrapure water was prepared using a MilliQ purification system (MilliporeSigma).

### Colon organoid culture and monolayer establishment

The colon monolayer model was derived from primary human colon organoids. Organoids from one donor (30 yr male patient for diverticulosis and diverticulitis, the normal appearing region of rectosigmoid) were obtained from Dr. O. Yilmaz lab at the Koch Institute/MIT. The medium used for maintaining organoids and monolayers include a base medium, organoid growth medium, seeding medium, and differentiation medium. The recipe for each medium is listed in **Table S8**.

The establishment and maintenance of the organoids were done according to the protocols previously described.^41,57^ In brief, organoids grown in Matrigel (growth factor reduced, phenol red free; Corning, 356231) droplets were passaged every seven days at a 1:3 split ratio. A medium change was performed on day four after passaging. To prepare the monolayer, organoids were collected at day 7 and pelleted by centrifugation (1000 g × 5min, 4°C). After that, organoid pellet was disrupted using Cell Recovery Solution (Corning, 354253; 1mL per 100 µL Matrigel). The resulting organoid suspension was then incubated on ice for 45-60 min, pelleted, and resuspended with 1 mL pre-warmed PBS without calcium and magnesium (PBS^-/-^, Gibco, 10010-023) containing 2.5mg/mL Trypsin (Sigma, T4549) and 0.45 mM EDTA (Ambion, AM9260G). The resuspended organoids were warmed up (37 °C water bath for 5 min) and then manually dissociated into single cells using a 1000-µL pipette with a bent tip. Trypsin was neutralized with 10% FBS in base medium. The cell suspension was then pelleted at 300 g × 5 min, 4°C. Finally, then cell pellet was resuspended in the seeding medium. Cell density and viability were determined using an automated cell counter (Invitrogen) and TrypanBlue. Before seeding, Transwells were coated with rat tail collagen I (Gibco, A10483-01, 50 µg mL^-1^ in PBS) for 1-2 hours in the incubator, then were washed with PBS right before adding the cells. For seeding, cells were diluted to a density of 600,000 cells per mL in seeding medium and seeded (500 µL) into the apical side of each 12-well collagen-coated Transwell (surface area: 1.12 cm^2^) and 1.5 mL cell-free seeding medium was added to the basolateral side. On day three after seeding, the monolayers were differentiated by switching to the antibiotic-free base medium on the apical side and differentiation medium on the basolateral side. After switching to differentiation medium, the monolayers were further cultured for four days (total seven days), with medium change on day five. On day seven after seeding (day four after differentiation), the monolayers were used for experiments.

### Bacteria culture and maintenance

*E. rectale* ATCC33656, *B. thetaiotaomicron* VPI-5482 were purchased from ATCC. *Faecalibacterium prausnitzii* DSM17677 was obtained from the Harvard Digestive Disease Center. The identity of all strains was confirmed using Sanger sequencing (see below). Bacteria from glycerol stock were plated in yeast casitone fatty acid (YCFA) agar (Anaerobe Systems, AS-675), 24-48 h after being cultured at 37 °C in the incubator inside the anaerobic chamber (Coy Laboratory), a colony was picked and cultured in Hungate tubes containing liquid YCFA medium (Anaerobe Systems, AS-680). Standard YFCA medium contains 33 mM acetate, 9 mM propionate and no butyrate.^12^ O_2_ in the anaerobic chamber was constantly removed by the Palladium Catalyst (Coy Laboratory, #6501050), which was renewed biweekly by incubating in the 90 °C oven for two days.

### Colon epithelial monolayer culture in the GuMI device

All components of the GuMI device **(Figure 1)** were sterilized by autoclave (121 °C, 45 min), except the pneumatic plates, oxygen probes, and probe controlling boxes, which were sterilized with ethylene oxide. Then the device was assembled under sterile conditions. Antibiotic-free base medium (1.5 mL; see **Table S8** for composition) was pipetted into the basal compartment. GuMI apical medium (110 mL), comprising filter-sterilized diluted YCFA medium (10% YFCA in PBS^+/+^) was added to the apical source reservoir on top of the GuMI device (total capacity 150 mL). The medium in the apical source reservoir was deoxygenized with 5% CO_2_, 95% N_2_ for 45-60 min before being introduced into the apical inlet through stainless steel tubing (**Figure 1**). After that, the apical inlet of the Transwell was temporally blocked with a 200-µl pipette tip to force the deoxygenized apical medium to flow out of the injection port, which was then sealed with an injection septum and a customized stainless-steel hollow screw. The pipette tips were then removed. The colon epithelial monolayers were transferred to each of the 6 basolateral reservoirs designed to accommodate standardized Transwells and the apical medium of the monolayers was replaced with the 10% diluted YCFA in PBS^+/+^. Then the entire basal plate was integrated with the apical plate using the lever (**Figure 1**). The system was primed 24 h in a cell culture incubator while the medium in the apical source reservoir was constantly purged with 5% CO_2_, 95% N_2_. The recirculation flow rate in the basal compartment was 5 µl/min and the apical flow rate was 10 µl/min. The effluent was cleared every 24 h with a 10-ml syringe (302995, BD Biosciences) throughout the experiments.

### Bacteria co-culture with colon epithelial monolayers

Colon epithelial monolayers were cultured in the GuMI device for 24 h before the addition of bacteria. The overnight grown bacterial cultures were diluted 1000 times with pre-reduced YCFA medium. After that, 0.8-1 ml of the diluted bacterial cells were slowly injected into the apical channel through the injection port (**Figure 1**) using a 1-ml syringe (309659, BD Biosciences) with a needle (305127, BD Biosciences). Before bacteria injection, the flow was paused to ensure the bacterial culture going through in one direction from the inlet and outlet of the apical channel. After one-hour settling of the bacterial cells, the flow was resumed in both apical and basolateral sides. At the end of the experiment, the whole device was transferred to a biosafety cabinet and the basal plate was carefully disassembled using the lever. The sealed Transwells were individually take off from the apical plate and placed onto a new 12-well plate. Immediately after that, the apical medium was collected using a 1-ml syringe with a short needle (305122, BD Biosciences), and then immediately injected into a 20-ml pre-reduced and autoclaved HDSP vial (C4020-201, Thermo Scientific) sealed with 20-mm Crimp Cap (95025-01-1S, MicroSolv). Then all the vials were transferred into an anaerobic chamber, where 10 µl of the apical medium was used for CFU counting on agar plates. The rest of the medium was transferred into a 1.5-ml polypropylene tube, where bacterial cells were pelleted in a microcentrifuge (14000 g × 5 min). The supernatant was transferred into a new tube. All samples were stored at -80 °C until further analysis.

The Transwells were washed with PBS^+/+^ (14040182, Thermo Scientific) twice in both apical and basolateral sides to completely remove cell-culture medium prior to bright field images and TEER measurement. After aspirating PBS, 350 μl of 1% 2-mercaptoethanol solution was added into the apical side, followed by incubation for 10 min at room temperature. One volume of 70% ethanol was then added and mixed homogeneously with pipetting. The mixture was collected and stored at -80 °C until further analysis.

### Immunofluorescence staining

Monolayers seeded in Transwells were washed with PBS^+/+^ and fixed in 4% formaldehyde for 10 minutes. The samples were then washed three times with PBS^+/+^ and permeabilized with 0.2% Triton-X for 10 minutes. After permeabilization, the wells were washed twice in PBS and blocked with Blockaid (Thermo Scientific B10710) for one hour. Primary antibodies diluted in Blockaid were incubated with samples overnight at 4°C. The following antibodies were used in the experiments at 1:200 dilution: anti-Muc2 (Abcam ab90007), anti-NHE3 (Novus, NBP1-82574). The samples were then washed three times in DPBS and incubated with secondary antibodies –Alexa Fluor 568 (1:200), phalloidin-Alexa Fluor 488 (1:20, ab176753) and DAPI (1:1000) diluted in Blockaid for one hour at room temperature. After washing the samples with PBS^+/+^ for three times, the monolayers were excised and mounted on coverslip using ProLong Gold antifade reagent (Thermo Fisher). Mounted samples were imaged with Zeiss LSM800 confocal microscope.^47^

The staining procedures were modified to preserve the bacterial cells attached to the colon epithelia (specifically for Figure 2h). Briefly, monolayers taken off from the platform were immediately fixed with 4% formaldehyde for 10 minutes following a very gentle sampling of the apical medium. The samples were then permeabilized with 0.2% Triton-X for 10 minutes. After permeabilization, the wells were washed once with PBS^+/+^ and immediately stained overnight with Phalloidin-iFluor 488 Reagent (ab176753-300TEST) and DAPI (1:1000) in Blockaid at 4 °C. After washing the samples with PBS^+/+^ for two times, the monolayers were excised, mounted, and imaged as described above.

### PCR and Sanger sequencing for bacteria

Bacteria identity was confirmed by Sanger sequencing by adapting the established protocol.^131^ Briefly, bacterial cells collected from the apical side of GuMI were collected and pelleted by centrifugation (12000 g × 5 min). The DNA was extracted using GeneElute bacterial DNA kit (NA2110, Sigma-Aldrich) by following the manufacturer protocol. Afterward, PCR was performed in triplicate to amplify 16s rDNA using DreamTaq Green PCR Master Mix (K1081, Thermo Fisher Scientific Inc.) with primers F8 (5’-AGTTTGATCCTGGCTCAG-3’) and 1492R (5’-TACGGYTACCTTGTTACGACTT-3’) by following the procedures described elsewhere.^55^ PCR products were purified using DNA purification solid-phase reversible immobilization magnetic beads (G95, Applied Biological Materials Inc.) and the purified products were sent out for Sanger sequencing (Genewiz Inc.).

### Multiplex cytokine/chemokine assays

The concentration of autocrine factors, cytokines, and chemokines in the apical media was measured using customized MILLIPLEX MAP assays, 47-plex human cytokine/TH17 panel (EMD Millipore) adapted from the previous protocol.^132^ Briefly, samples were measured at multiple dilutions to ensure the measurements were within the linear dynamic range of the assay. We reconstituted the protein standard in the same media and serially diluted the protein stock to generate a 7-point standard curve. Assays were run on a Bio-Plex 3D Suspension Array System (Bio-Rad Laboratories, Inc.). Data were collected using the xPONENT for FLEXMAP 3D software, version 4.2 (Luminex Corporation, Austin, TX, USA). The concentration of each analyte was determined from a standard curve that was generated by fitting a 5-parameter logistic regression of mean fluorescence on known concentrations of each analyte (Bio-Plex Manager software).

### Mucin measurement

Mucin measurement is performed according to the method previously reported^133^ and adapted to the 96-well plate. Briefly, Dye stock solution contains 1% (w/v) alcian blue (A5268, Sigma-Aldrich) and 0.098% (v/v) H_2_SO_4_. Before diluting to dye working solution, dye stock solution was syringe filtered (0.22 µm PVDF, Cat No 09-720-3, Fisher Scientific) to remove the particles. Dye working solution containing 0.25% Triton X-100, 0.098% H_2_SO_4_, and 5% dye stock solution was syringe filtered. Ten microliters of reaction solution (0.147% v/v H_2_SO_4_, 0.375% Triton X-100, and 4 M guanidine HCl) was added to each 10 µl of standard series (0, 4.84, 9.68, 19.37, 38.75, 67.5, 125, and 250 µg/mL) in bacterial medium (10% YCFA in PBS^+/+^) along with 100 µl of dye working solution. After centrifugation for 20 min at 2400 g, the supernatant was removed by reverting the 96-well plate to the bed of tissue papers. Then, an 8M guanidine HCl solution was added to dissolve the pellet. The absorbance at 600 nm was recorded and mucin concentration was determined from a calibration curve. Technical duplicates were included for all the samples and standards.

### Liquid chromatography-mass spectrometry (LC-MS) analysis

Short chain fatty acids, butyrate, propionate, and acetate, were quantified by LC-MS using the derivatization method reported elsewhere.^134^ Briefly, SCFA in the samples were derivatized by mixing with 20 µl sample with 20 µl 3-nitrophenylhydrazine, and 20 µl 120 mM *N*-(3-dimethylaminopropyl)-*N*′-ethylcarbodiimide HCl. The reaction mixture was incubated at 37 °C for 30 min before being diluted 20 times with 9:1-water:acetonitrile mixture. The diluted samples were then filtered with 0.22 µm PTFE membrane. The filtrate was collected and immediately injected for LC-MS analysis or store at -20 °C until analysis within a week. An Agilent 1200 HPLC system (Agilent) coupled to a Q-TOF mass spectrometry (Agilent) and an electrospray ionization (ESI) source was used. Chromatographic separation was carried out using a Phenonemix NH_2_ column (4.6 ×150 mm, 4.6 µm). Analytes were eluted with mobile phase A (water with acetic acid) and B (acetonitrile with acetic acid) at 0.35 ml/min with the following gradient program: 0-2 min 35% B; 15.5-17 min 98% B; 17.5-25 35% B. The elute was ionized in negative mode with following parameters: gas temperature 300 °C, drying gas 8 l/min, nebulizer 30 psi, sheath as temperature 350 °C, sheath gas flow 12 l/min, MS TOF fragmentor at 70 V, skimmer 65 V. Targeted m/z for acetate, propionate, and butyrate, and isotope labeled butyrate were 194.0571, 208.0728, 222.0884, and 229.1324, respectively. The detection limit for all analytes was 0.08 mM.

### RNA extraction and quality control

Prior to extraction, the cell lysate in 1% 2-mercaptoethanol solution was mixed with one volume 350 µl of 70% ethanol and pipetted to a homogeneous mixture. Then total RNA was extracted using PureLink RNA mini kit (ThermoFisher, 12183020) by following the manufacture protocol, except treating samples with PureLink DNase (ThermoFisher, 12185010) during one of the wash steps to remove DNA. The total RNA was analyzed by Bioanalyzer in BioMciroCenter at MIT. All the RNA samples passed the QC with RNA quality number (RQN) 9.8-10.0.

### Library preparation and Illumina sequencing

The library preparation, sequencing, and analysis were carried out in BioMicro Center at MIT. Briefly, 50ng of RNA was confirmed for quality using the Agilent Fragment Analyzer and 50ng of material was polyA selected using NEBNext® Poly(A) mRNA Magnetic Isolation Module (E7490) modified to include two rounds of polyA binding and 10-minute incubations. cDNA was generated using the NEB Ultra II directional kit (E7760) following the manufacturer’s protocol using 15 cycles of PCR and a 0.9X SPRI clean. The resulting libraries were quality assessed using the Agilent Fragment Analyzer and quantified by qPCR prior to being sequenced on the Illumina HiSeq2000. The 40nt single-end reads with an average depth of 5 million reads per sample were sequenced for all conditions.

### RNA-seq data analysis

#### Differential gene expression

To guarantee data quality for downstream analyses, QC was performed by MIT BioMicro Center in-house QC software (script upon request) to monitor sequencing error rate, GC bias, sequence count, mapping rate, contamination, CDS percentage, UTR percentage, intro percentage, intergenic percentage, exon to intron ratio, 5’ to 3’ ratio, sense to antisense ratio, rRNA percentage, sequence complexity, and number of genes detected. The single-end sequences were mapped to GRCh38 reference sequence by STAR 2.5.3a using the following parameters: --outFilterType BySJout: keep only those reads that contain junctions that passed filtering into SJ.out.tab; --outFilterMultimapNmax 20: alignment will be output only if it has fewer mismatches than 20; --alignSJoverhangMin 8: minimum overhang for unannotated junctions 8bp; --alignSJDBoverhangMin 1: minimum overhang for annotated junctions 1bp; --outFilterMismatchNmax 999: maximum number of mismatches 999. Large number switches off this filter; --alignIntronMin 10: minimum intron size: genomic gap is considered intron if its length>=10, otherwise it is considered Deletion; --alignIntronMax 1000000: maximum intron size 1000000 bp; --alignMatesGapMax 1000000: maximum gap between two mate 1000000bp; --outSAMtype BAM SortedByCoordinate: output sorted by coordinate Aligned.sortedByCoord.out.bamfile, similar to samtools sort command; --quantMode TranscriptomeSAM: outputs alignments translated into transcript coordinates in the Aligned.toTranscriptome.out.bam file (in addition to alignments in genomic coordinates in Aligned.*.sam/bamfiles).

The sorted bam files were further indexed using samtools 1.3. Gene expression was estimated by calculating the gene level raw read counts, FPKM, and TPM using rsem 1.3.0 rsem-calculate-expression. Due to the strand specificity of the library, only reverse strand was counted using –forward-prob 0 option. In addition, the command –calc-pme option was applied to run collapsed Gibbs sampler for posterior mean estimation. The raw counts, log2(FPKM+1), and log2(TPM+1) from all samples were merged into 3 corresponding tables using MIT BioMicro Center in-house tools (script available upon request). Differential expression was performed using DESeq2 1.10.1 based on gene level raw counts. Pair-wise comparisons were performed across two conditions: NB versus S, and FP versus NB. To reduce the burder for multi-testing correction, genes with no expression in any samples during comparison were filtered away. up-regulated and the down-regulated genes were further selected by MIT BioMicro Center in-house tools. Up-regulation is defined as: baseMean>10, log2FoldChange>0.5, and adjusted p <0.05. Down-regulation is defined as: baseMean>10, log2FoldChange<-0.5, and adjusted p <0.05.

#### Gene set enrichment analysis

Gene set enrichment analysis were performed by GSEA 4.0.3.^64^ log2(TPM+1) of the expressed genes in each comparison were project to compiled gene sets from literature or databases that have curated gene sets (Hallmarks and MSigDB). The significantly enriched gene set is defined as nominal p < 0.05, and FDR q-value <0.05.

#### Ingenuity Pathway Analysis (IPA)

The identified differentially expressed genes was uploaded in IPA to identify the networks and upstream regulators that are most significant in GuMI-NB vs Static or GuMI-FP vs GuMI-NB by Causal Networks Analysis and Upstream Regulators Analysis.^135^

### Reverse transcription quantitative polymerase chain reaction (RT-qPCR)

RT-qPCR was performed to quantify gene expression. Briefly, the mRNA was converted to cDNA using the High-Capacity RNA-to-cDNA Kit (Thermo Fisher Scientific, 4387406). TaqMan Fast Advanced Master Mix (Thermo Fisher Scientific, 4444557) and TaqMan probe (**Table S9**) were mixed in MicroAmp EnduraPlate Optical 96-well fast clear reaction plate with barcode (Thermo Fisher Scientific, 4483485) according to manufacture protocol. TaqMan probes used in this study are available in **Table S9**.

### Transepithelial electrical resistance measurement

EndOhm-12 chamber with an EVOM2 meter (World Precision Instruments) was used to measure the transepithelial electrical resistance (TEER) values.

### Computational Simulation

A finite element method was performed using COMSOL Multiphysics 5.4 (COMSOL Inc.). Two modules (laminar flow fluid dynamics and transport of diluted species) were coupled to compute the flow distribution and the profile of oxygen concentration inside the apical and basal compartment of the MPS. The 3D CAD files of the MPS were drawn using Solidworks (Dassaut Systèmes®) and exported into COMSOL. Only half of the 3D MPS was simulated given the plane of symmetry along the feeding axis. First, the steady state solution for the computational fluid dynamics (CFD) was calculated using the Navier-Stokes equation assuming incompressible fluid. As boundary condition for the CFD, the interfaces between the medium and walls of the MPS (solid walls as well as on the Transwell membrane on both sides) were set as no slip conditions. The interface between the medium and air was set as a stationary free interface with intrinsic surface tension. The linear flow rates (μL min^-1^) at both inlets (basal in recirculation and apical for feeding) were set according to experimental measurements, and were assumed to be fully developed. The two outlets (basal for recirculation and apical for effluent) were set as outflows for the simulation (degree of freedom for pressure). To simulate the transport of oxygen molecules, a diffusion/convection model was applied using Fick’s 2^nd^ law. The oxygen consumption rate (OCR) was calculated based on published OCR from intestinal epithelial cells. The total number of epithelial cells on the Transwell was estimated and the thickness of the tissue was measured using confocal microscopy. The flux of oxygen through the membrane of the Transwell was allowed through passive diffusion using the manufacturers porosity. The inlet on the apical side was set at a fixed oxygen concentration (O_2_ deprived medium). On the basal compartment, the inlet and outlet coming in and out of the pump are constrained to the same oxygen concentration using a continuity condition. The standard mesh was applied, and all simulations were performed with the assumption that the system is at 37 °C, in an air composed of 5% CO_2_ and with 100% humidity.

A first simulation was performed as time-dependent to characterize how fast the system reaches equilibrium (steady state oxygen delivery). A second study was performed at steady state to fully characterize oxygen and shear stress distribution in the MPS. Parameters used in the simulations are provided in **Table S1**.

An estimate of the purge time of tiny bubbles entrained during placement of the Transwell was carried out as follows: Assuming a 50-µl air bubble underneath the upper insert into the apical compartment that forms the upper wall of the apical flow channel, with 0.4 cm^2^ of contact area with liquid medium. The half-life of diffusion from the entrained air bubble to the liquid medium is about 5.6 h. At flow rate 10 μl min^-1^, the speed of purging is 0.183 × 10^−8^ mol min^-1^. As the concentration of oxygen in the bubble is equal to m/v = 0.2 × 1.01 × 10^5^/8.314/310 = 7.84 mol m^-3^. The time required for purging the medium is C × V/ Speed = 7.84 mol m^-3^ × 50 × 10^−9^ m^3^/ (0.183 × 10^−8^ mol/min) = 3.5 h. Diffusion is longer than purging. Therefore, diffusion from the air bubble to apical medium is the decisive factor for reaching equilibrium. Generally, it takes about five half-lives to replace 99% of oxygen in the air bubble, (about 18.5 h). Together with 2.5 h to purge liquid medium, the total time is very close to the experimentally observed time for the oxygen reading to reach equilibrium.

## Ethic statement

Colon organoids used in this study were established by the Yilmaz lab at the Koch Institute/MIT (HC2978 – 30 yr male patient for diverticulosis and diverticulitis, the normal appearing region of rectosigmoid sample used). Endoscopic tissue biopsies were collected from the ascending colon of de-identified individuals at either Massachusetts General Hospital upon the donors informed consent. Methods were carried out in accordance with the Koch Institute Institutional Review Board Committee as well as the Massachusetts Institute of Technology Committee on the use of humans as experimental subjects.

## Acknowledgment

This study was supported by the NIH R01EB021908 and the Boehringer Ingelheim SHINE Program. This work was supported in part by the National Institute of Environmental Health Sciences of the NIH under award P30-ES002109. We are grateful for S. Levine and D. Ma at MIT BioMicro Center for their help with RNA-seq data analysis. We thank B. Joughin for helpful discussion, and S. Manalis and the MIT 20.330 students for inspiration to analyze diffusion and reaction in the mucus layer.

## Conflict of Interest

A provisional patent was filed.

