## Supplemental Figures for "Primary human colonic mucosal barrier crosstalk with super oxygen-sensitive *Faecalibacterium prausnitzii* in continuous culture"

**Contribution:**

Hardware design, testing, and troubleshooting: YJH, JYY, DLT, LGG, JK, BB, GR, KB, DC, SJH, JCK

Bacterial culture: JZ, YJH, TM, CAV

Experimental design: JZ, YJH, LGG, DLT

Experiment execution: JZ, YJH, CW, KS, JK

Computational modeling: PS

Cell culture: YJH, CW, KS, VHG, JZ, DTB, GE, OY

Data analysis: JZ, YJH, LGG, DLT

Figures and Tables configuration: JZ, YJH

Writing: JZ, YJH, LGG, DLT

Supervision: LGG, DLT, RC

All authors commented and approved the manuscript.

**Supplementary Methods**

**Supplemental Method A:** Characteristic Diffusion time for SCFA in Apical Compartment

The characteristic transient diffusion time is τ_D_ ~ L^2^/D_AB_, where D_AB_ is the diffusion coefficient. For molecules in the molecular weight range of glucose and SCFA diffusing in culture media or loose mucus, DAB ~0.5-1 x 10^-5^ cm^2^/s) at 37F.

Hence, for L = 0.3 cm (the height of the apical compartment above the epithelial layer, assuming the mucus layer is negligible)

is τ_D_ ~ L^2^/D_AB_ ~ (0.3 cm)^2^/(5 x 10^-6^ cm^2^/s) = 18,000 s = 5 hr. Thus, butyrate produced in the mucus layer will not saturate the apical flow during its ~0.5 hr transit through the apical compartment.

**Supplemental Method B:** Glucose diffusion is not limiting the fermentation in *F. prausnitzii* layer

Assumptions:

To estimate how the ~150 µm thickness of the *F. prausnitzii* layer that appears to be embedded in a loose mucus layer above the epithelial barrier (Fig 2), we presume the following:

(i) the microbes are evenly distributed in a mucus layer of L = 150 µm thickness

(ii) no convection in the mucus layer

(iii) culture medium flowing over the top of the mucus layer contains a bulk concentration of glucose C_bulk_ = 2.8 mM (2.8 × 10^-6^ mole/cm^3^) as the main precursor for butyrate fermentation

(iv) glucose diffuses into the mucus microbial mass from the bulk in a one dimensional manner and is consumed at a zero-order volumetric rate, which is approximately equal to the production rate of butyrate; this assumption does not directly account for glucose consumed for bacterial growth except through the carbon stoichiometry differences (glucose = 6, butyrate = 4).

(v) the bulk apical medium is the only source of butyrate

Thiele modulus – dimensionless group reflecting ratio of reaction:diffusion

A mass balance on 1 diffusion and with zero reaction in the bacterial mass yields the dimensionless parameter called the Thiele modulus, F^2^.^136^

F^2^ = Q_g_L^2^/(D_g_C_bulk_)

here Q_g_ is the volumetric butyrate production rate (measured) and D_g_ is the diffusion coefficient for glucose in the bacterial mass. For values of F^2^ >2, the reaction is strongly diffusion limited as the substrate concentration drops to zero at the far edge of the diffusion/reaction path when F^2^ = 2. For metabolic reactions, further analysis could be conducted using first-order or non-linear (e.g. Michaelis-Menten) rate expressions; the Thiele modulus approach is an order-of-magnitude assessment of diffusion limitations.

Data used

(i) The diffusion coefficient of glucose is 1 × 10^-6^ cm^2^/s.

(ii) Bulk concentration of glucose = 2.8 mM

(iii) Production rate of butyrate = 19 + 1.6 + 0.2 µmol = 22 µmol in 48 hr (data in Table 1).

With L = 150 µm thick layer of mucus/bacteria, V_bacteria_ = 1.1 cm^2^ × 0.015 cm = 0.0165 cm^3^

Q_g_ = 2.2 × 10^-5^ mol/0.0165 cm^3^/48 hr/3600s/hr = 7.7 × 10^-9^ mole/cm^3^/s

Analysis

F^2^ = Q_g_L^2^/(D_g_C_bulk_)

= (7.7 × 10^-9^ mole/cm^3^/s)(0.015cm)^2^/(1 × 10^-5^ cm^2^/s)/(2.8 × 10^-6^ mol/cm^3^)

= 0.06

For these conditions, the fermentation reaction is not limited by diffusion of glucose.

**Supplemental Method C:** Exemplified estimations of the concentration of cytokine, chemokine and growth factor at cell surface in GuMI-NB: IL-8 and TGF-α

Assumptions

To estimate the cell surface concentrations of secreted or shed apical proteins in the presence of apical flow, we presume the following:

(i) the flux of protein from the apical surface into the bulk is constant and occurs by 1-D diffusion in the z direction from the surface, and can be estimated by dividing the total accumulation in the static condition over 48 hr by the culture surface area (1.1 cm^2^); the similarity of flux between static and flow conditions is supported by the lack of significant differences at the transcriptional level.

(ii) the consumption of protein by autocrine processes (receptor binding and internalization) is negligible compared to diffusion of protein into the bulk.

(iii) A layer of mucus of thickness L = 150 µm covers the apical surface, and no convection occurs in this region.

(iv) the concentration of autocrine factors in the bulk flow regime is zero.

Flux Balance and Surface Concentration Equation

With these assumptions, and designating the diffusion coefficient of protein in mucus as D_pm_, the protein concentration at the surface as C_surface_, invocation of Fick’s law of diffusion with the boundary condition of C_bulk_ = 0 @ x = L, yields the following flux balance:

Flux = (D_pm_/L)(C_surface_ – 0)

Rearrange to yield the cell surface concentration

C_surface_ = Flux(L/ D_pm_)

Data used

(i) Production rate of IL8 = (7171 pg/mL)(0.5 mL)/48hr/3600s•hr^-1^/1.1cm2 = 0.019 pg/cm^2^/s

Production rate of TGF-a = (61.8 pg/mL)(0.5 mL)/48hr/3600s•hr^-1^/1.1cm2 = 0.00016 pg/cm^2^/s

(ii) Molecular weights: IL-8 = 8.4 kDa, TGF-a = 5.5 kDa

(iii) Diffusion coefficient D_pm_ ~ 5 × 10^-6^ cm^2^/s

Estimated concentrations at the cell surface:

IL8: C_surface_ = 0.019 pg/cm^2^/s [(0.015 cm/5 × 10^-6^ cm^2^/s) = 57 pg/mL = 6.9 pM

TGF-a: C_surface_ = 0.00016 pg/cm^2^/s [(0.015 cm/5 × 10^-6^ cm^2^/s) = 0.49 pg/mL = 0.09 pM

**Supplementary Tables in the separated Excel File**

**Table S1.** Parameters used for the simulation of flow pattern and oxygen distribution

**Table S2.** Absolute amount of SCFA under Static, GuMI-NB, and GuMI-FP conditions.

**Table S3.** The list of genes that are significantly changed in GuMI-NB compared with Static cells. Corresponding to Figure 3a.

**Table S4.** The list of genes that are significantly changed in GuMI-FP compared with GuMI-NB. Corresponding to Figure 3b.

**Table S5.** The list of core genes in the cell differentiation gene set enrichment analysis in Figure 4c.

**Table S6.** IPA revealing upstream regulators in GuMI-NB compared with Static cells.

**Table S7.** IPA revealing upstream regulators in GuMI-FP vs GuMI-NB cells.

**Table S8.** Composition of all media used for colon organoids and monolayers.

**Table S9.** TaqMan probes used for RT-qPCR.

**Supplementary figures**

**Figure S1.** Oxygen and shear stress distribution in GuMI physiome platform. (a) oxygen distribution in the apical and basolateral sides of colon epithelia after reaching equilibrium. (b) Time course of oxygen in apical side. (c) distribution of shear stress across the monolayer (top-down view). (d) Shear stress distribution. (e) side view and (f) top-down view of the simulation for the flow streamline

**Figure S2.** Bright field images of the monolayer cultured under Static (a), GuMI (b), and GuMI-FP (c). bar scale = 300 µm. (d) Concentration of short chain fatty acids in effluent from in GuMI-NB and GuMI-FP after 48 h of co-culturing.

**Figure S3.** Overview of the effects of GuMI and *F. prausnitzii* on the gene expression of colon epithelial cells. (**a**) volcano plot of the genes in GuMI vs Static, the significantly changed genes are highlighted in green; total n is the number of genes with TPM > 0 in all samples, sig. n is the number of genes that are significantly changed (adj. p<0.05, log2FoldChange>0.5). (**b**) volcano plot of the significantly changed genes in GuMI+FP vs GuMI, the significantly changed genes are highlighted in red; (**c**) Overlap on the genes changed by GuMI and *F. prausnitzii*. The number in the circle indicates the number of genes changed uniquely by only either condition or by both conditions. The chromosol distribution of the changed genes in GuMI vs Static(**d**) and GuMI+FP vs GuMI (**e**). (**f**) Heatmap of the core genes that are significantly under-represented in NB, and the function of these genes in DNA replication machinery for pre-initiation, initiation, elongation, and maturation of DNA. MCM: DNA helicase family minichromosomal maintenance protein complex; CDC: cell division cycle protein; GINS2, GINS Complex Subunit 2; POLA1-2, DNA polymerase alpha 1 and 2; POLD3, DNA polymerase delta 3; PCNA, proliferating cell nuclear antigen; RFC, replication factor C subunit; ORC, origin recognition complex subunit; LIG1: DNA ligase.

**Figure S4.** Concentration of cytokine/chemokines in the apical media and the expression of corresponding genes in the epithelial cells at Static, GuMI-NB, and GuMI-FP. (**a**) Heatmap of the core genes that are significantly under-represented in NB, and the function of these genes in DNA replication machinery for pre-initiation, initiation, elongation, and maturation of DNA. MCM: DNA helicase family minichromosomal maintenance protein complex; CDC: cell division cycle protein; GINS2, GINS Complex Subunit 2; POLA1-2, DNA polymerase alpha 1 and 2; POLD3, DNA polymerase delta 3; PCNA, proliferating cell nuclear antigen; RFC, replication factor C subunit; ORC, origin recognition complex subunit; LIG1: DNA ligase. (**b**) Ten factors with autocrine signaling properties in colon epithelia were detected at concentrations of 3 – 20 pM. The mean transcript, gene expression fold change and significance in GuMI-NB vs Static, GuMI-FP vs GuMI-NB were displayed beneath GuMI-NB and GuMI-FP, respectively. FC: fold change; adj. p: adjusted p value; ND: no transcript detected in RNA-seq. Note the unit of protein concentration is ng/L in the figures. (**c**) Twelve factors were detected at concentrations of 0.3-2 pM range. FC: fold change; adj. p: adjusted p value; ND: no transcript detected in RNA-seq. Note the unit of protein concentration is ng/L in the figures. (**d**) and (d) 20 factors were undetectable above background. FC: fold change; adj. p: adjusted p value; ND: no transcript detected in RNA-seq. Note the unit of protein concentration is ng/L in the figures. (**e**) IL-1β and IL-33 were not detected at protein level but were significantly changed in GuMI-NB comparing to Static and in presence of *F. prausnitzii*. FC: fold change; adj. p: adjusted p value; ND: no transcript detected in RNA-seq. Note the unit of protein concentration is ng/L in the figures. (**f**) No significant changes of HIF1A negative regulator genes EGLN1, EGLN2, EGLN3, HIF1AN in GuMI-NB vs Static. NB/ST: GuMI-NB/Static. (**g**) Changes of TLR3 and TLR4 and their downstream genes in GuMI-FP vs GuMI-NB.

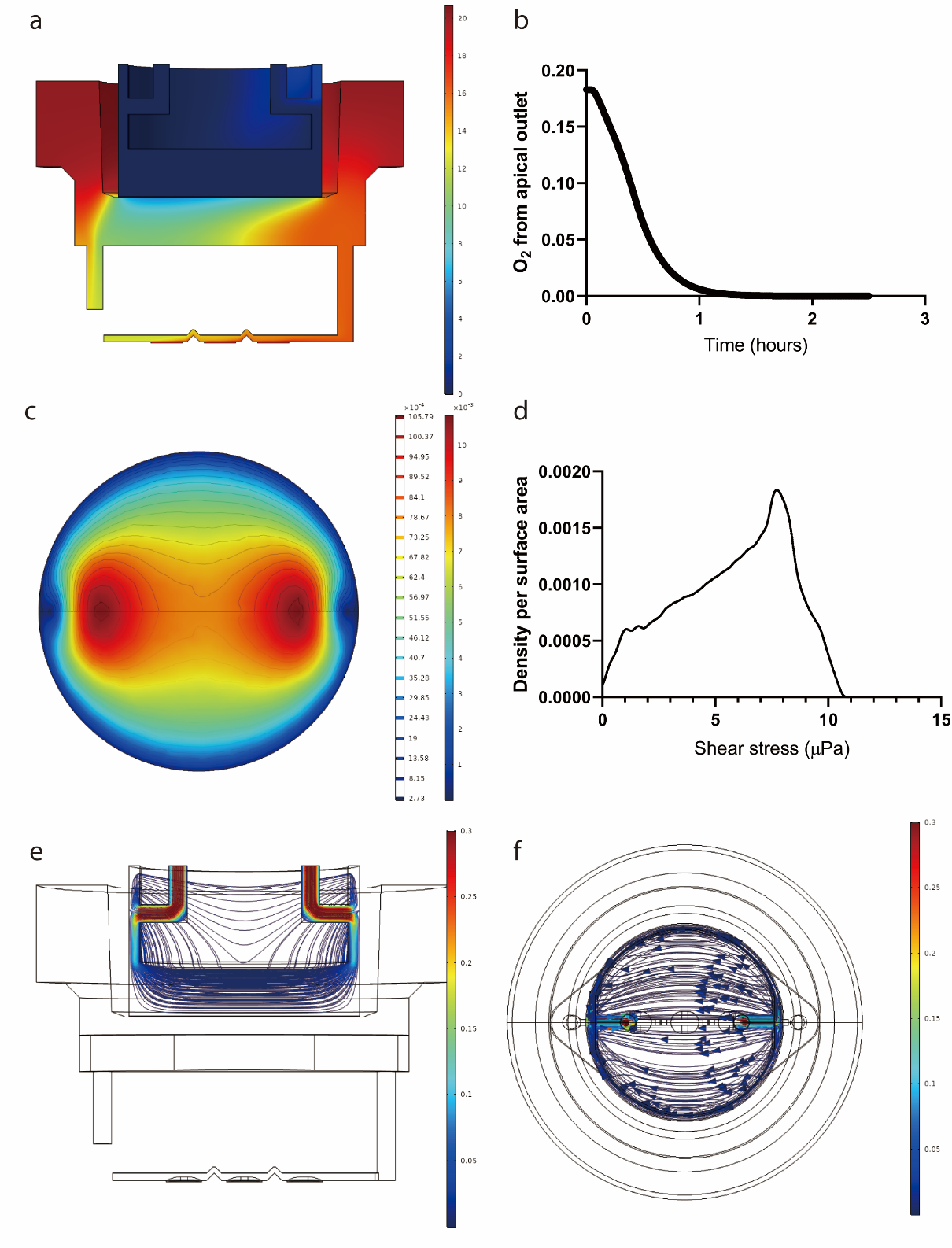

**Figure S1.** Oxygen and shear stress distribution in GuMI physiome platform. (a) oxygen distribution in the apical and basolateral sides of colon epithelia after reaching equilibrium. Concentrations on the basolateral side were measured in pilot experiments. (b) Time course of oxygen in apical side. (c) distribution of shear stress across the monolayer (top-down view). (d) Shear stress distribution. (e) side view and (f) top-down view of the simulation for the flow streamline

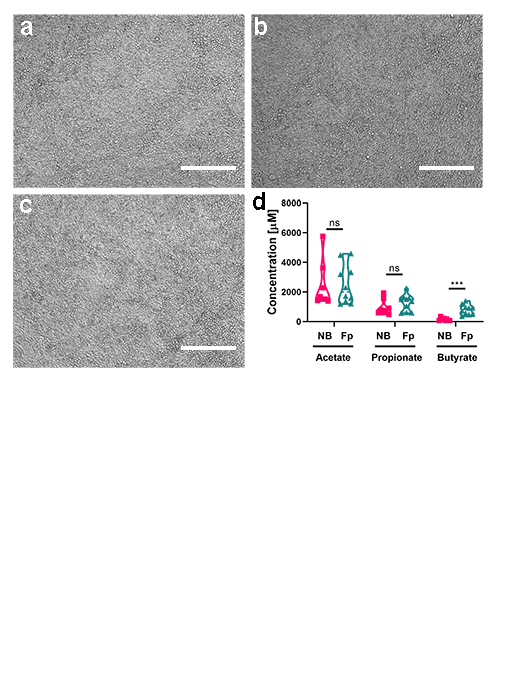

**Figure S2.** Bright field images of the monolayer cultured under Static (a), GuMI (b), and GuMI-FP (c). bar scale = 300 µm. (d) Concentration of short chain fatty acids in effluent from in GuMI-NB and GuMI-FP after 48 h of co-culturing.

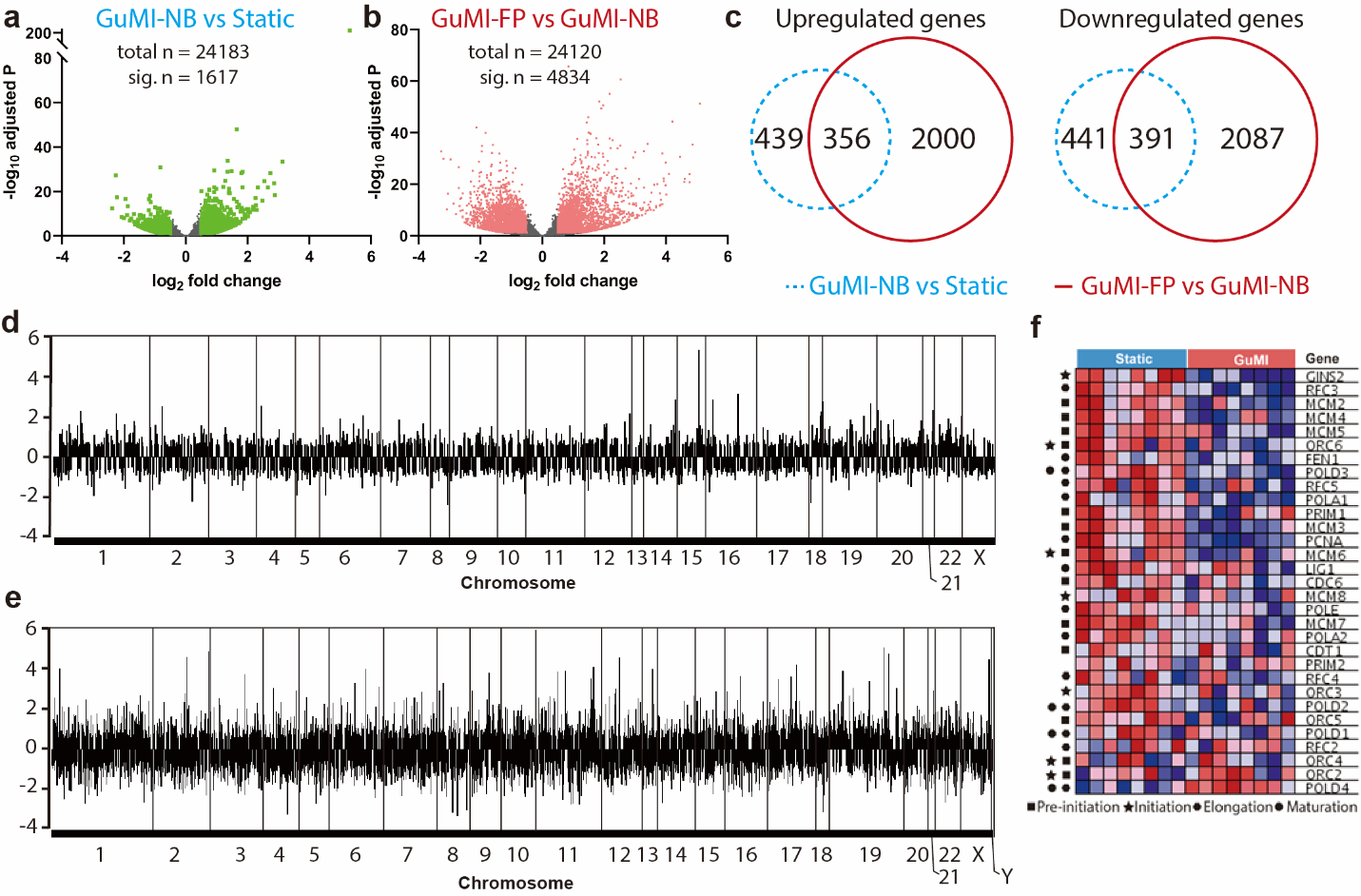

**Figure S3.** Overview of the effects of GuMI and *F. prausnitzii* on the gene expression of colon epithelial cells. (**a**) volcano plot of the genes in GuMI vs Static, the significantly changed genes are highlighted in green; total n is the number of genes with TPM > 0 in all samples, sig. n is the number of genes that are significantly changed (adj. p<0.05, log2FoldChange>0.5). (**b**) volcano plot of the significantly changed genes in GuMI+FP vs GuMI, the significantly changed genes are highlighted in red; (**c**) Overlap on the genes changed by GuMI and *F. prausnitzii*. The number in the circle indicates the number of genes changed uniquely by only either condition or by both conditions. The chromosol distribution of the changed genes in GuMI vs Static(**d**) and GuMI+FP vs GuMI (**e**). (**f**) Heatmap of the core genes that are significantly under-represented in NB, and the function of these genes in DNA replication machinery for pre-initiation, initiation, elongation, and maturation of DNA. MCM: DNA helicase family minichromosomal maintenance protein complex; CDC: cell division cycle protein; GINS2, GINS Complex Subunit 2; POLA1-2, DNA polymerase alpha 1 and 2; POLD3, DNA polymerase delta 3; PCNA, proliferating cell nuclear antigen; RFC, replication factor C subunit; ORC, origin recognition complex subunit; LIG1: DNA ligase.

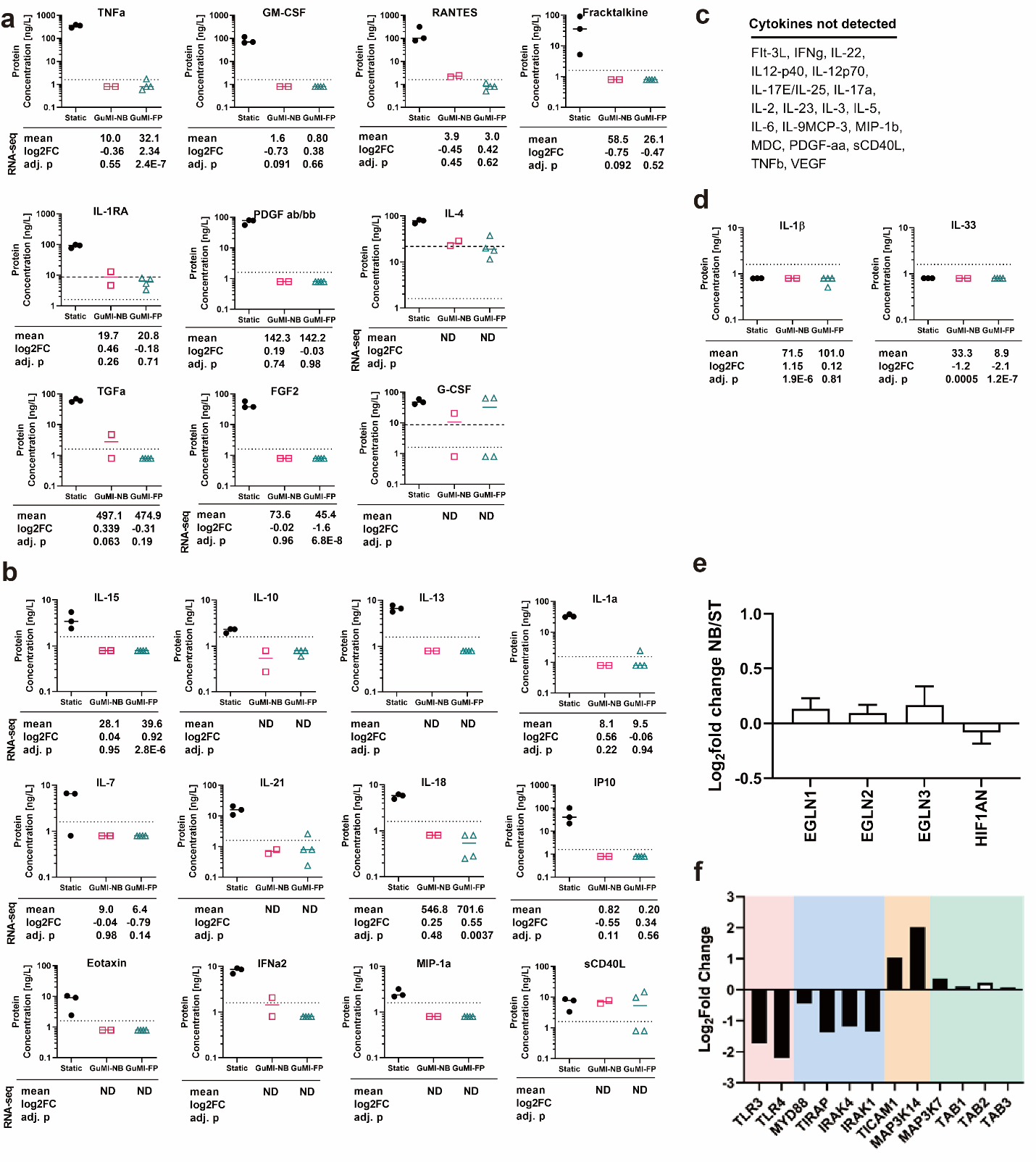

**Figure S4.** Concentration of cytokine/chemokines in the apical media and the expression of corresponding genes in the epithelial cells at Static, GuMI-NB, and GuMI-FP. (**a**) Ten factors with autocrine signaling properties in colon epithelia were detected at concentrations of 3 – 20 pM. The mean transcript, gene expression fold change and significance in GuMI-NB vs Static, GuMI-FP vs GuMI-NB were displayed beneath GuMI-NB and GuMI-FP, respectively. FC: fold change; adj. p: adjusted p value; ND: no transcript detected in RNA-seq. Note the unit of protein concentration is ng/L in the figures. (**b**) Twelve factors were detected at concentrations of 0.3-2 pM range. FC: fold change; adj. p: adjusted p value; ND: no transcript detected in RNA-seq. Note the unit of protein concentration is ng/L in the figures. (**c**) and (d) 20 factors were undetectable above background. FC: fold change; adj. p: adjusted p value; ND: no transcript detected in RNA-seq. Note the unit of protein concentration is ng/L in the figures. (**d**) IL-1β and IL-33 were not detected at protein level but were significantly changed in GuMI-NB comparing to Static and in presence of *F. prausnitzii*. FC: fold change; adj. p: adjusted p value; ND: no transcript detected in RNA-seq. Note the unit of protein concentration is ng/L in the figures. (**e**) No significant changes of HIF1A negative regulator genes EGLN1, EGLN2, EGLN3, HIF1AN in GuMI-NB vs Static. NB/ST: GuMI-NB/Static. (**f**) Changes of TLR3 and TLR4 and their downstream genes in GuMI-FP vs GuMI-NB.
